# Lymphoma B cells remodel bone marrow stromal cell organization and function to induce a supportive cancer-associated fibroblast network

**DOI:** 10.1101/2023.09.26.559605

**Authors:** Elise Dessauge, Baptiste Brauge, Simon Léonard, David Roulois, Céline Monvoisin, Thomas Lejeune, Jérôme Destin, Florence Jouan, Francisco Llamas-Gutierrez, Frédéric Mourcin, Karin Tarte

**Affiliations:** UMR 1236, Univ Rennes, INSERM, Etablissement Français du Sang, Equipe Labellisée Ligue, F-35000, Rennes, France; Laboratoire d’Anatomie Pathologique, Centre Hospitalier de Rennes, Rennes, France; SITI laboratory, CHU Rennes, Etablissement Français du Sang, F-35000, Rennes, France

## Abstract

Bone marrow (BM) involvement is a common feature of lymphomas deriving from germinal-center B cells and is associated with a bad prognosis. In particular, follicular lymphoma (FL) infiltrates the BM in 70% of cases, in association with a remodeling of surrounding tumor microenvironment. Analysis of *in vitro*-expanded FL mesenchymal stromal cells (MSCs) revealed an extensive alteration of BM stromal cell phenotypic, transcriptomic, and functional profiles. However, the mechanisms supporting the direct interplay between lymphoma B cells and their permissive stromal niche *in situ* have not been yet identified. In the current work, we identified in the BM milieu of FL patients a deregulation of soluble and extracellular matrix (ECM) components reflecting inflammation and ectopic differentiation into lymphoid-like stromal cells. We reproduced the same alterations in a murine model of lymphoma B-cell xenograft where a scRNAseq approach identified LepR^pos^ MSCs as specifically and progressively reprogramed by tumor B-cell invasion. Analysis of FL BM collected before and after treatment confirmed that BM niche was partly dependent on the continuous contact with tumor B cells. Altogether, this work shed new lights on the kinetic and mechanisms of BM stromal niche reshaping in B-cell lymphoma.

## Introduction

Follicular lymphoma (FL) and diffuse large B-cell lymphoma (DLBCL) are the two most common B-cell lymphomas^1–3^. FL is an indolent disease, characterized by the accumulation of germinal center (GC)-derived B cells with a disseminated pattern, infiltrating the bone marrow (BM) in 50-70% of cases at diagnosis^3^. Compared to lymph node (LN) FL B cells, BM FL B cells remain organized as nodular follicle-like aggregates but are characterized by a lower cytological grade, a reduced proliferation, a modified phenotype with loss of CD10 expression, and a specific transcriptomic profile reflecting reduced metabolic activity^4–6^. Furthermore, BM is a niche for long-lived lymphoma precursor cells, as highlighted by the synchronous development of a clonally-related FL pair in donor and recipient of allogeneic BM transplantation^7^. DLBCL is a group of GC/post-GC aggressive lymphomas with a high degree of molecular heterogeneity and a BM involvement in 10% to 25% of patients^8^. FL and GC-derived DLBCL (GCB-DLBCL), in contrast to non-GC derived B-cell lymphomas, share a preferential paratrabecular niche in invaded BM^9^. Of note, FL transforms into DLBCL, usually of GCB-type, at a rate of 3% per year^10,11^. BM involvement is an adverse prognostic factor in both FL and DLBCL and contributes to lymphoma risk-stratification scores^8,12–14^. The BM microenvironment has long been thought to provide a supportive niche for tumor B cells throughout lymphoma development, from pre-tumoral stage to overt lymphoma, relapse, and/or transformation^15^, but the composition and function of BM lymphoma-supportive niches and how they acquire their protumoral activity remain elusive.

FL is the paradigm of a neoplasia strongly dependent on a complex and dynamic tumor microenvironment (TME), including reprogrammed CD4^pos^ T cells, myeloid cells, and lymphoid stromal cells (LSCs), that are closely related to the normal GC niche but exhibit specific phenotypic, transcriptomic, and functional features^3,16^. FL-infiltrating LSCs are extensively remodeled *in situ* within FL LN, showing deregulated production of chemokines and extracellular matrix (ECM) components, thus forming an organized network of heterogeneous cancer-associated fibroblasts (CAFs)^17^. Similarly, mesenchymal stromal cells (MSCs) obtained from FL BM display a specific transcriptomic profile associating an ectopic LSC-like signature with upregulation of *CCL2* and *IL-8*^18,19^. In accordance, FL BM-MSCs have both a greater capacity to directly support FL B-cell survival and to recruit protumoral monocytes and neutrophils compared to BM-MSCs obtained from healthy donors (HD). Importantly, all of these factors are inducible *in vitro* by coculture of HD BM-MSCs with malignant B cells, suggesting a direct role for FL B cells in the priming of BM FL-CAFs, particularly through their production of tumor-necrosis factor α (TNF) and lymphotoxin α1β2 (LT), the two key non-redundant drivers of LSC differentiation and maintenance^18–20^. Conversely, CXCL12 is also specifically upregulated in FL LN and BM stromal cells but is not induced by the contact with FL B cells *in vitro*^21^. Both CD4^pos^ T cell-derived IL-4 and extracellular vesicles (EVs) produced by FL B cells have been proposed as inducers of CXCL12 upregulation in FL BM-MSCs^4,21^. FL-derived EVs may play a role in the initial induction of tumor-permissive niche within BM, in the absence of infiltration by tumor cells and immune TME. Importantly, while purified LN FL-CAFs have been extensively characterized by scRNAseq approaches^22^, data on BM FL-CAFs have been obtained using long-term *in vitro* expanded cells derived from BM aspirates, without taking into account the functional heterogeneity of BM stromal cells. Yet, seminal papers have proposed a molecular atlas of BM stromal cells, providing clues as to how different stromal cell subtypes can support the different steps of normal B-cell hematopoiesis^23,24^. In DLBCL, despite convincing data revealing LSC remodeling within invaded LN, associated with alteration of chemokine and ECM profiles^25^, and *in vitro* studies suggesting a role for B-cell/stromal cell crosstalk within invaded BM^26,27^, BM DLBCL-CAFs have never been explored.

In the current study, we used BM samples from lymphoma patients and a model of lymphoma B-cell xenograft in immunocompromised mice to decipher the reprogramming of BM stromal cells in lymphoma-invaded BM. By analyzing stromal cell heterogeneity using scRNAseq and immunohistofluorescence approaches, we identified LepR^pos^ MSCs as the major cell compartment affected by tumor B-cell invasion and described a deregulation of chemokine production and ECM organization that was directly driven by B-cell growth. This work sheds new light on the kinetics of BM stromal niche remodeling in lymphoma and on the role of the continuous crosstalk between B cells and BM-stromal cells in the polarization of the BM TME.

## Results

### Characterization of the medullary stromal niche in BM-invaded FL patients

We previously demonstrated that *in vitro*-expanded FL BM-MSCs overexpress the inflammatory chemokines CCL2 and IL-8 compared to HD BM-MSCs and proposed local FL B-cell-derived TNF as an inducing mechanism^18,19^. Moreover, we identified FL B-cell-derived TNF and transforming growth factor β (TGFβ) as involved in the upregulation of lymphoid chemokine expression, in particular CCL19, in LN FL-CAFs^17^. In order to characterize BM stromal cell reprogramming into LSC-like cells in FL patients *in situ* we carried out BM plasma dosages of chemokines and cytokines associated with LSC niche, comparing early and later stage FL patients with invaded BM to age-matched HD (Figure 1A). Both CXCL13, mainly produced by LN follicular dendritic cells (FDCs), and CCL19, mainly produced by LN fibroblastic reticular cells (FRCs), were found significantly upregulated in FL BM plasma compared to HD BM plasma (P<.0001), highlighting the induction of a functional ectopic LSC niche. CXCL13 and CCL19 levels were correlated in FL BM plasma (P<.0001), suggesting a common regulating mechanism. Additionally, the level of CXCL13 and CCL19 in invaded BM were higher in high-tumor burden (n=19) FL than in low-tumor burden (n=12) FL (P<.001 and P<.05; respectively). Looking for driving factors, we next identified a significant upregulation of both TNF and TGFβ1 in FL BM plasma (P<.0001) and TNF, unlike TGFβ1, was found strongly correlated to CCL19 and CXCL13 expression (Supplemental Figure S1). Finally, the inflammatory soluble factors CXCL10 and IL-6, but also the immunosuppressive cytokine IL-10 were all upregulated in FL BM, underlying, besides LSC commitment, a large deregulation of the BM milieu in the presence of FL B cells. Since LN FL-CAFs are characterized by a huge deregulation of chemokines and ECM^17^, we next performed staining of FL-invaded BM biopsies for ECM components, focusing on Collagen VI, as a marker strongly expressed by LN FRC^28^. In FL-infiltrated BM, Collagen VI network was amplified and reorganized compared with HD BM (Figure 1B). Altogether, these results reinforce the demonstration of local TME remodeling in FL-invaded BM and argue for a direct impact of tumor B cells on BM stroma reprogramming into LSC-like cells.

**Figure 1.**
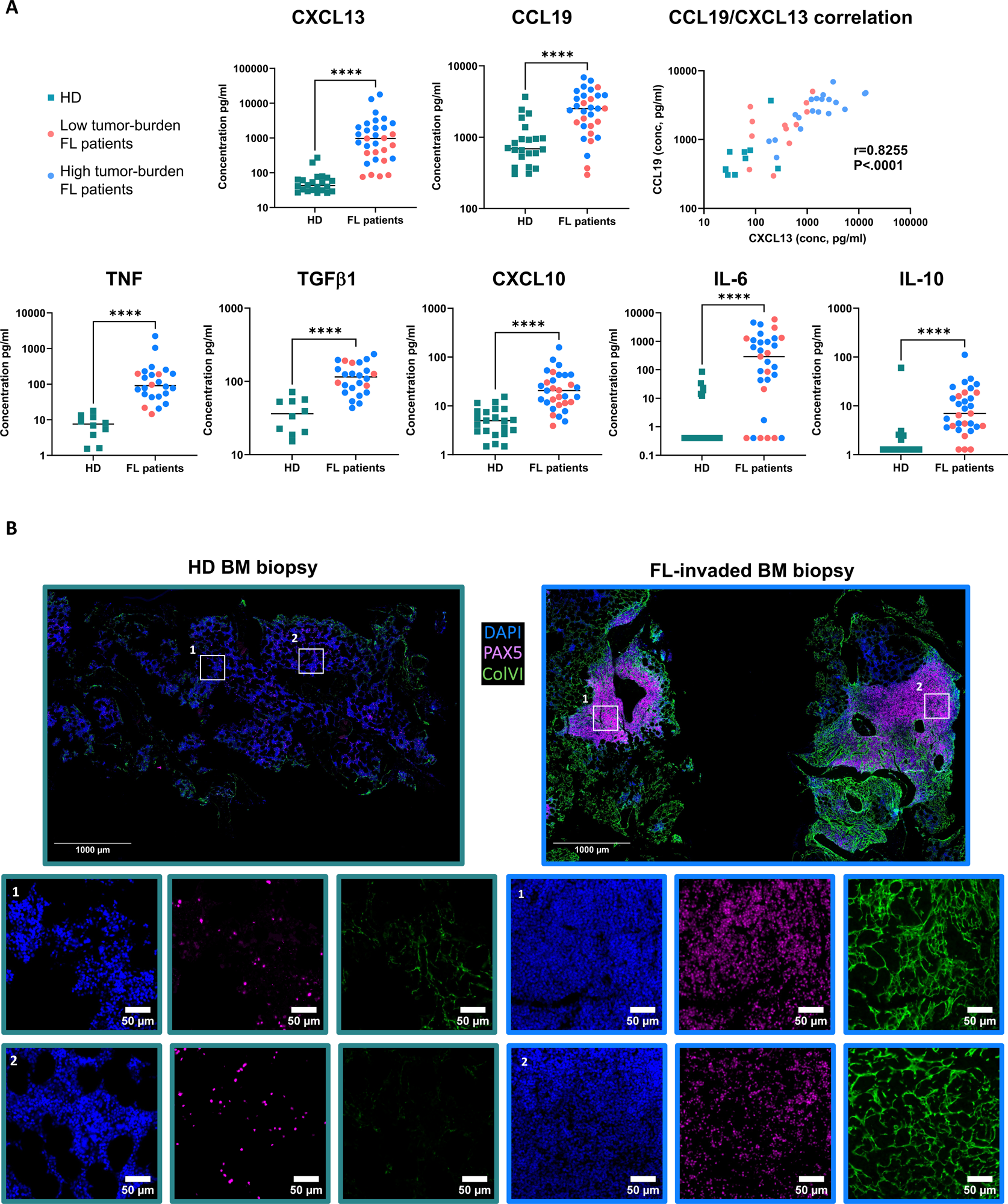
Characterization of the medullary niche composition in BM-invaded FL patients. (A) Chemokine and cytokine levels were measured in BM plasma from FL patients (n=29, except for TNF and TGF-β n=24) and HD (n=22, except for TNF and TGF-β n=10) using Luminex assay. Green squares represent HD, blue dots represent high tumor-burden FL patients and pink dots represent low tumor-burden FL patients. **** P< .0001. (B) Immunofluorescence on BM sections of healthy donors (HD) and FL patient for PAX5 (magenta) and Collagen VI (green). Nuclei were counterstained with DAPI. Scale bar, 1000 µm. Boxes indicate areas magnified in upper panels where scale bars represent 50µm.

### Intrafemoral xenograft model reproduces lymphoma BM niche

To decipher the direct crosstalk between lymphoma B cells and the heterogeneous BM stromal cell compartment, we developed an intrafemoral xenograft model in Rag^-/-^γc^-/-^ mice and evaluated two GC-derived lymphoma B-cell lines: the GCB-DLBCL cell line OCI-Ly-19 and the FL cell line DOHH2. At 0.5×10^6^ cells/mice, the rate of engraftment was of 100% for both cell line but with a significantly different kinetic, as highlighted by a median day of euthanasia of 25 for OCI-Ly-19- and 42 for DOHH2-grafted mice (P<.001; Supplemental Figure S2A). In addition, whereas OCI-Ly-19 graft produced a rapid progression and a systemic dissemination in all mouse tissues (data not shown), quantification of hCD20^pos^ cells by flow cytometry and detection of Luc-DOHH2 cells using bioluminescence imaging revealed an initial engraftment of DOHH2 in the grafted femur followed by a specific expansion in the contralateral femur (Figure 2A). This slower tumor progression and preferential BM tropism, together with the previous demonstration that DOHH2 is dependent on the presence of stromal cells to grow in a 3D spheroid model^29^, support the use of DOHH2 intrafemoral xenograft model to mimic B-cell lymphoma BM localization. Moreover, to capture the kinetic of B cell/stromal cell crosstalk, we defined day 40 (D40) as the later evaluable time point (Gr.Late) and day 19 (D19) as the time point where BM involvement was 10-fold lower than at D40 (Gr.Early) (Supplemental Figure S2B).

**Figure 2.**
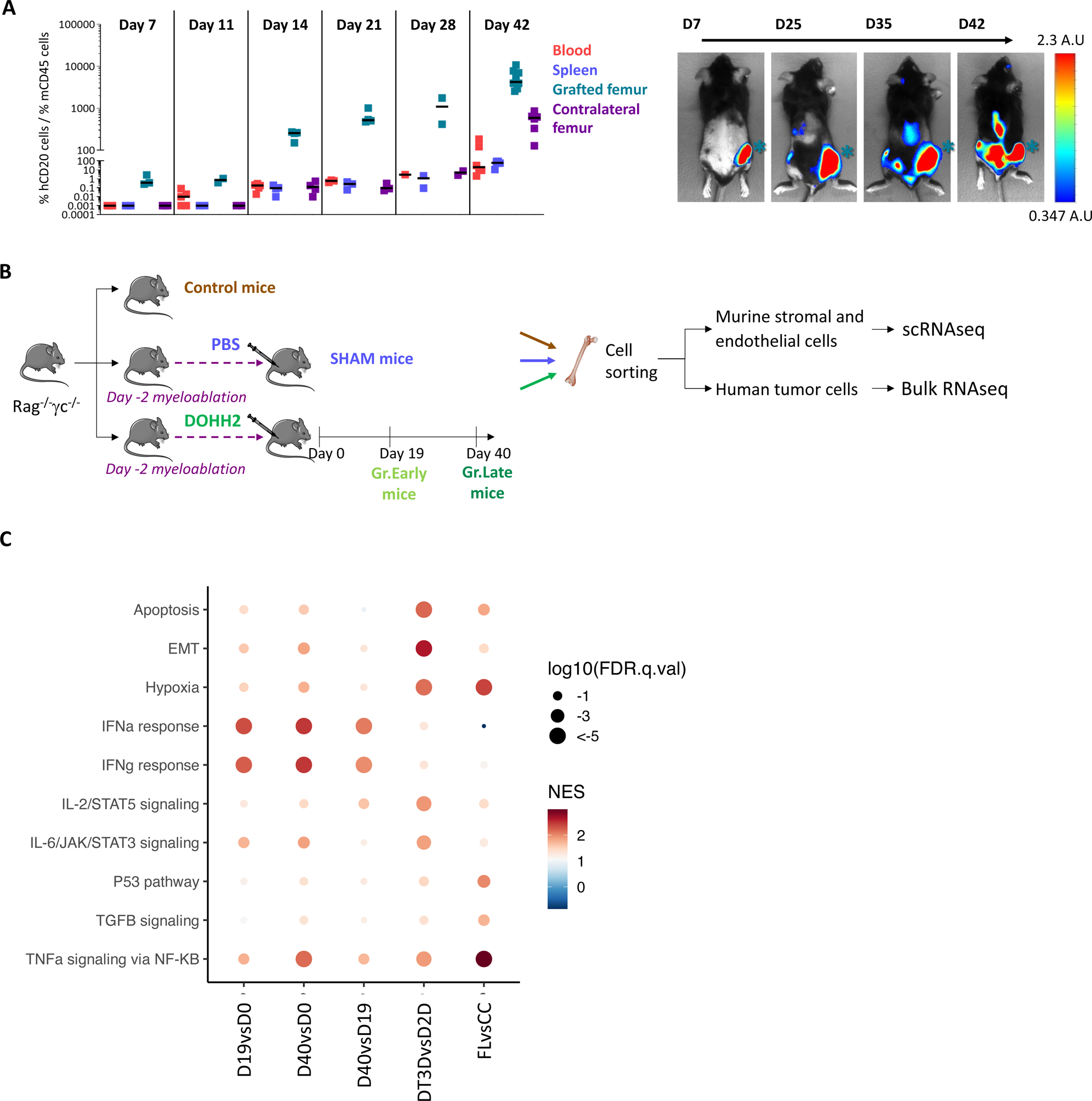
Establishment and characterization of a B cell/stroma cell crosstalk mouse model. (A) Tumor cell dissemination. Evolution over time of the proportion of viable human tumor cells (DAPI^neg^hCD20^pos^) compared to viable murine CD45^pos^ cells in blood (red), spleen (blue), and BM (grafted femur in green and contralateral femur in purple) of Rag^-/-^γc^-/-^ mice transplanted with 0.5×10^6^ DOHH2 cells by intrafemoral route (left panel). Luciferase imaging of representative mouse from 7 to 42 days post-engraftment (right panel). (B) Experimental design of the B cell and stromal cell transcriptomic characterization in Rag^-/-^ γc^-/-^ mice untreated, injected with PBS (Sham), or grafted with DOHH2. (C) Enrichment for pathways upregulated in DOHH2 recovered at the late time point (D40) and/or at the early time point (D19) and relevant to lymphoma biology were studied using GSEA in published data of DOHH2 co-cultured with tonsil stromal cells and ECM in 3D spheroids (DT3D) *versus* classical 2D DOHH2 culture (D2D) (Lamaison *et al*, Blood Adv 2020) and of primary FL B cells (FL) versus centrocytes (CC) (Desmots *et al*., Clin Cancer Res 2019). Circle colors depict the normalized enrichment score (NES) and circle sizes the false discovery rate (FDR).

Based on these data, we designed a kinetic study of non-hematopoietic BM cell reprogramming in response to lymphoma B-cell xenograft (Figure 2B). For that purpose, we sorted viable Lin^neg^CD31^pos^SCA-1^pos^ endothelial cells and Lin^neg^CD51^pos^CD200^pos^ stromal cells (Supplemental Figure 2C) before any manipulation (control mice), in PBS-grafted mice at D40 (Sham mice), and in DOHH2-grafted mice at D19 (Gr.Early mice) and D40 (Gr.Late mice), and analyzed them by scRNASeq. Moreover, we studied by bulk RNAseq the gene expression profile (GEP) of DOHH2 at D19 and D40 compared to that of DOHH2 before transplantation (D0) and determined differentially expressed genes (DEGs) between these 3 conditions using DESeq2 analysis (n=3; adjusted P-value < .05) (Supplemental Table S1). We next applied gene set enrichment analysis (GSEA) to these gene lists and highlighted progressively upregulated pathways relevant to lymphoma biology, including TNF, interferon (IFN), TGFβ, and IL-6 signaling and hypoxia (Figure 2C). Interestingly, similarly reprogrammed pathways were found in published datasets of DOHH2 encapsulated in 3D spheroids in the presence of tonsil stromal cells and ECM compared to classical 2D culture, a model found to reproduce *in vitro* FL B cell/stromal cell crosstalk^29^, but also in primary purified FL B cells compared to normal centrocytes^30^, suggesting that our *in vivo* xenograft model is relevant to FL biology and will be useful to capture B cell/stromal cell interactions.

### Lymphoma B cells reprogram LepR^pos^ MSCs in invaded BM

Our scRNA-seq dataset contained stromal and endothelial cells from control mice, Sham mice, Gr.Early mice, and Gr.Late mice and corresponded, after quality control analysis, to a total of 15,590 cells organized in 15 clusters after unbiased UMAP clustering (Figure 3A, left). We used a previously published atlas identifying 17 clusters of non-hematopoietic bone and BM cells in steady-state C57BL/6 mice^24^ to generate corresponding specific scores and identify the main cell subsets in our dataset of Rag^-/-^γc^-/-^ mice (Figure 3A, right). We identified five clusters of chondrocytes and fibroblasts (clusters 8, 12, 9, 5, and 14), a cluster of pericytes (cluster 11), four clusters of sinusoidal and arteriolar/arterial endothelial cells (clusters 2, 3, 10 and 7), two clusters of osteolineage cells (OLCs, clusters 6 and 13), and three clusters of Leptin receptor (LepR)^pos^ MSCs (clusters 4, 0, and 1). Cluster 15 corresponded to cycling cells. We next assessed the repartition of the different experimental conditions in each cluster (Figure 3B). As determined by a Chi^2^ test, the most significantly unbalanced clusters between control, Sham, Gr.Early, and Gr.Late mice were clusters 4, 0, and 1, corresponding to the three LepR^pos^ MSC clusters. Cluster 4 was almost exclusively composed of LepR^pos^ MSCs from control mice, cluster 0 was strongly enriched for Sham mouse LepR^pos^ MSCs, and cluster 1 was mainly composed of LepR^pos^ MSCs from grafted mice. Interestingly, the Lepr-MSC signature obtained from steady-state C57BL/6 mice was less enriched in cluster 1 than in clusters 0 and 4, despite high *Lepr* expression (Supplemental Figure S3A-B). These data identified DOHH2-grafted LepR^pos^ MSCs as displaying a specific GEP, distinct from that of LepR^pos^ MSCs from immunocompetent mice with a normal B-cell hematopoiesis. Based on these observations, we decided to focus our study on LepR^pos^ MSCs.

**Figure 3.**
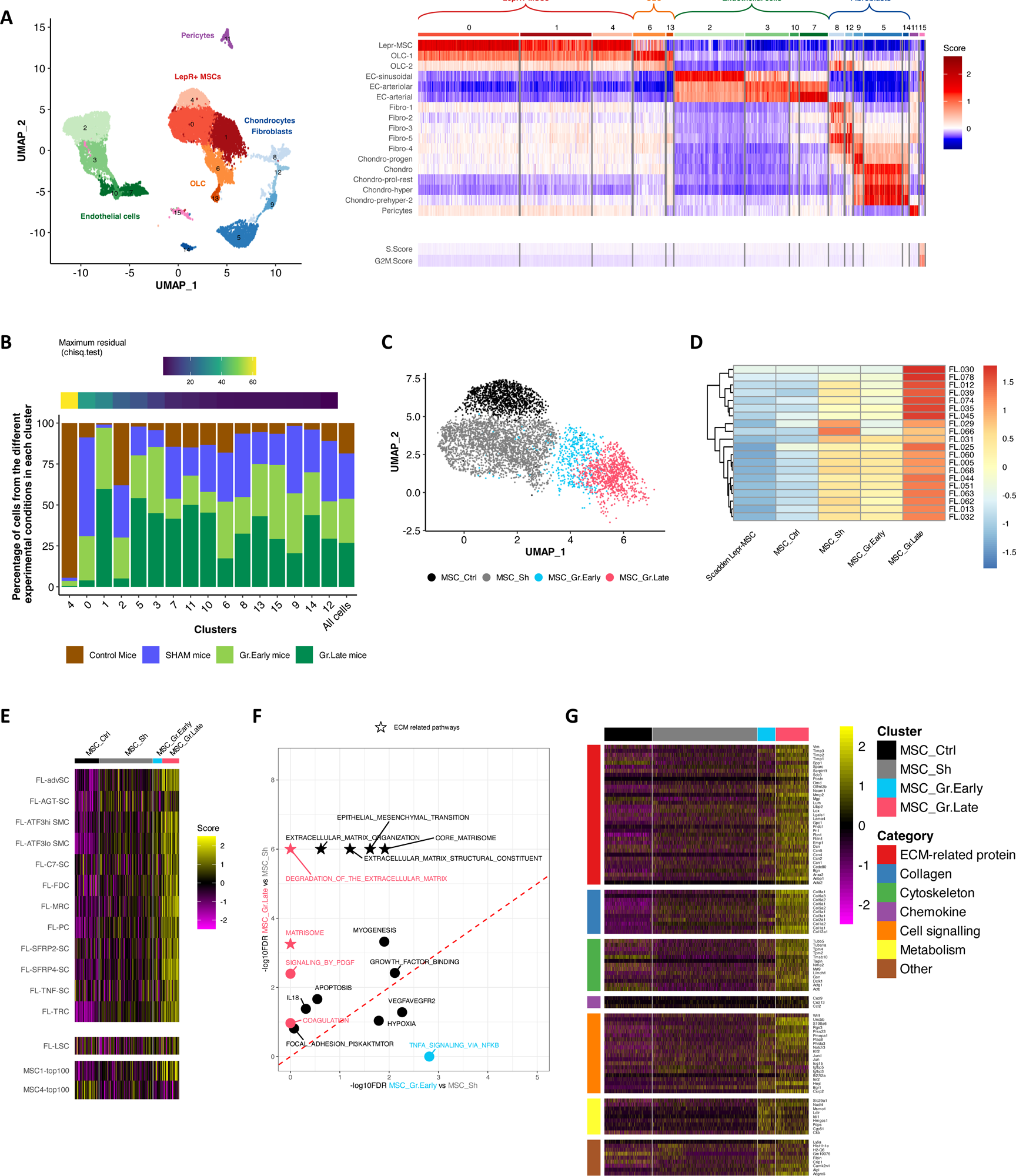
Lymphoma B cells drive LepR^pos^ MSC transcriptional reprogramming. (A) UMAP plot of the scRNAseq data from endothelial and stromal cells of Rag^-/-^γc^-/-^ mice untreated, injected with PBS, or grafted with DOHH2 and euthanized at early and late time-points (left panel). Heatmap of the enrichment of mouse non-hematopoietic cell signature scores as previously defined by scRNAseq in steady-state C57BL/6 mice (Baryawno *et al*., Cell 2019) (right panel). (B) Diagram representing, for each cluster, the proportion of cells coming from the four conditions (control, Sham, Gr.Early, and Gr.Late). The last bar of the histogram represents the distribution of all cells between the different experimental conditions. The balance between experimental conditions in each cluster was evaluated using a Chi^2^ test. (C) UMAP plot of the subclustering of the 3 most unbalanced clusters (4, 0, and 1) after analysis of differential abundance using the DA-seq algorithm. Four LepR^pos^ MSC sub clusters were identified: MSC_Ctrl, MSC_Sh, MSC_Gr.Early, and MSC_Gr.Late. (D) Heatmap generated with CellphoneDB repository, representing the number of significative bi-directional interactions between LepR^pos^ clusters (MSC_Sh, MSC_Gr.Early, MSC_Gr.Late, MSC_Ctrl from the current study and Lepr-MSC from Baryawno *et al.,* Cell 2019) and human primary FL B cells (Han G *et al*., Blood Cancer Discov 2022). (E) Heatmap of gene expression values from the human FL LN stromal cell subcluster signatures (published in Abe *et al.*, Nat Cell Biol 2022; upper), the human FL-LSC signature (published in Mourcin *et al*. Immunity 2021; middle), and the BM MSC clusters obtained from HD and MM patients (published in de Jong *et al.* Nat Immunol 2021; lower), relative to the LepR^pos^ MSC clusters identified by DA-seq. (F) Scatter plot of the significantly upregulated pathways in MSC_Gr.Late vs MSC_Sh (y-axis) and MSC_Gr.Early vs MSC_Sh (x-axis) as determined by GSEA analysis (Hallmarks, Canonical pathways and Gene Ontology databases). Pathways specific of each comparison are color-coded: red for MSC_Gr.Late versus MSC_Sh, blue for MSC_Gr.Early versus MSC_Sh, and black ones for common pathways. ECM-related pathways are indicated by a star. (G) Heatmap of the 100 genes upregulated in MSC_Gr.Late and/or MSC_Gr.Early.

To finely characterize the condition-dependent heterogeneity of LepR^pos^ MSCs and distinguish early from late time-points, we next used the differentially abundant-seq (DA-seq) multiscale approach designed to delineate cell subpopulations with the most significant discrepancy between different biological states, that are not captured by unsupervised clustering-based methods^31^. Applying this strategy on our LepR^pos^ MSCs, we obtained four clusters of DA populations, one enriched for LepR^pos^ MSCs from control mice (MSC_Ctrl), one enriched for LepR^pos^ MSC from Sham mice (MSC_Sh) and two clusters enriched for LepR^pos^ MSC from Gr.Early or Gr.Late mice (named MSC_Gr.Early and MSC_Gr.Late; respectively) (Figure 3C). Comparison of the DEGs (adjusted P-value < .05, absolute value of Log2FC>0.25) between the different conditions revealed that the MSC_Gr.Early vs MSC_Sh signature was largely included within the MSC_Gr.Late vs MSC_Sh signature (147/182 genes, 81%) whereas 311 out of the 646 genes of the MSC_Gr.Late vs MSC_Sh signature (48%) were common with MSC_Gr.Late vs MSC_Gr.Early signature (Supplemental Figure S3C and Supplemental Table S2). These data suggested that MSC_Gr.Early displayed an intermediate phenotypic state between MSC_Sh and MSC_Gr.Late.

The functional impact of this transcriptomic reprogramming on the capacity of BM-MSCs to interact with malignant B cells was studied using the CellPhoneDB repository, making it possible to comprehensively study biologically relevant ligand-receptor interactions in order to infer cell-cell communication networks^32^. For that purpose, we used public scRNASeq data obtained from 20 purified primary FL B cells^33^ and estimated the number of bidirectional interactions between these malignant B cells and LepR^pos^ MSCs, including our four clusters (MSC_Ctr, MSC_Sh, MSC_Gr.Early, MSC_Gr.Late) and LepR^pos^ MSCs from steady-state immunocompetent mice (Scadden_Lepr-MSC^24^) (Figure 3D). Interestingly, a higher number of cell interactions were predicted to occur between FL B cells and MSC_Gr.Late, thus reinforcing the hypothesis that MSC_Gr.Late acquired a specific capacity to interact with lymphoma B cells and that this lymphoma-induced MSC GEP was different from the GEP resulting from the crosstalk between MSCs and differentiating normal B cells in a normal BM niche. We next assessed whether our murine MSC clusters were differentially enriched for signatures of human FL-CAFs. Both whole FL-LSC signature identified by comparing LSCs sorted from FL LN *vs* reactive inflamed secondary lymphoid organs^17^ and FL-LSC subset signatures generated using a scRNAseq approach on stromal cells from FL LN *vs* normal LN^22^ were enriched in MSC_Gr.Late compared to MSC_Gr.Early, MSC_Sh, and MSC_Ctrl (Figure 3E and Supplemental Figure S3D). In order to explore a mature B-cell malignancy with primary BM involvement, we also analyzed scRNAseq data of human BM stromal cells purified from multiple myeloma (MM) patients and HD^34^ (Figure 3E and Supplemental Figure S3E). Whereas MSC_Gr.Late showed an enrichment for the MSC1 cluster signature, corresponding mainly to MM BM-MSCs, MSC_Ctrl were significantly enriched for the MSC4 cluster signature, enriched in HD BM-MSCs.

Finally, we explored the molecular pathways deregulated in BM LepR^pos^ MSCs in contact with lymphoma B cells. For that purpose, we applied GSEA to our MSC_Gr.Early vs MSC_Sh and MSC_Gr.Late vs MSC_Sh signatures (Figure 3F). This analysis highlighted a large set of signatures related to ECM as deregulated in both early and late stages, with a stronger upregulation in MSC_Gr.Late. When focusing on the 100 most upregulated genes in MSC_Gr.Early and/or MSC_Gr.Late, we observed in particular a huge enrichment for ECM-related genes, including *Sparc, Postn, Mmp2, Lox, Fn1,* or *Acta2*, cytoskeleton-related genes, including *Actb*, *Tpm2,* and *Tpm4*, collagens, including *Col6a1, Col6a2,* and *Col6a3*, and chemokines, including *Cxcl9, Cxcl13 and Ccl2* (Figure 3G). These signatures were enriched in MSC_Gr.Late compared to MSC_Gr.Early. Conversely, genes related to cell signalling, including *Jun, Heyl, Isg15*, or *Igfbp3/5*, and cholesterol metabolism, such as *Ldlr, Fdsp*, or *Cyp51*, were upregulated early in MSC. We then took advantage of our previous study comparing the GEP of *ex vivo* expanded BM-MSCs obtained from histologically invaded vs non-invaded FL patients^18^. Genes upregulated in MSC_Gr.Early and/or MSC_Gr.Late were found more upregulated in BM-MSCs from FL-invaded BM compared to BM-MSCs from FL non-invaded BM (Supplemental Figure S3F). These results add further evidence toward a change of the secretory profile and a substantial ECM remodeling in lymphoma-primed BM MSCs.

Importantly, we confirmed the deregulation of inflammatory chemokines, including Ccl2, Il-6, and Cxcl10 in the BM of DOHH2-grafted mice, as reported in FL patients^18^ (Figure 4A). Moreover, immunohistofluorescence on BM sections of Sham and Gr.Late mouse femurs showed that B-cell infiltration was associated with an induction of collagen VI and Sparc, a matricellular protein previously reported as expressed in BM infiltrated by GC-derived lymphomas^9^, in stromal cells (Figure 4B and Supplemental Figure S4A). To evaluate the relevance of this ECM remodeling, we investigated the enrichment of previously published DLBCL matrisome signatures^35^ in the four DA-seq generated clusters of LepR^pos^ MSCs. LepR^pos^ MSCs from DOHH2-grafted mice, especially MSC_Gr.Late, were found significantly enriched for lymphoma matrisome secreted factor, ECM-affiliated protein, collagen, and ECM glycoprotein signatures (Figure 4C and Supplemental Figure S4B). Similarly, MSC_Gr.Late were significantly enriched for myofibroblastic CAF (myCAF) unlike inflammatory CAF (iCAF) signatures identified previously in solid cancers^46^, and more precisely to TGF-β secreting and ECM organizing myCAF signatures^37^, suggesting some common features of ECM deregulation in lymphoma and solid cancers.

**Figure 4.**
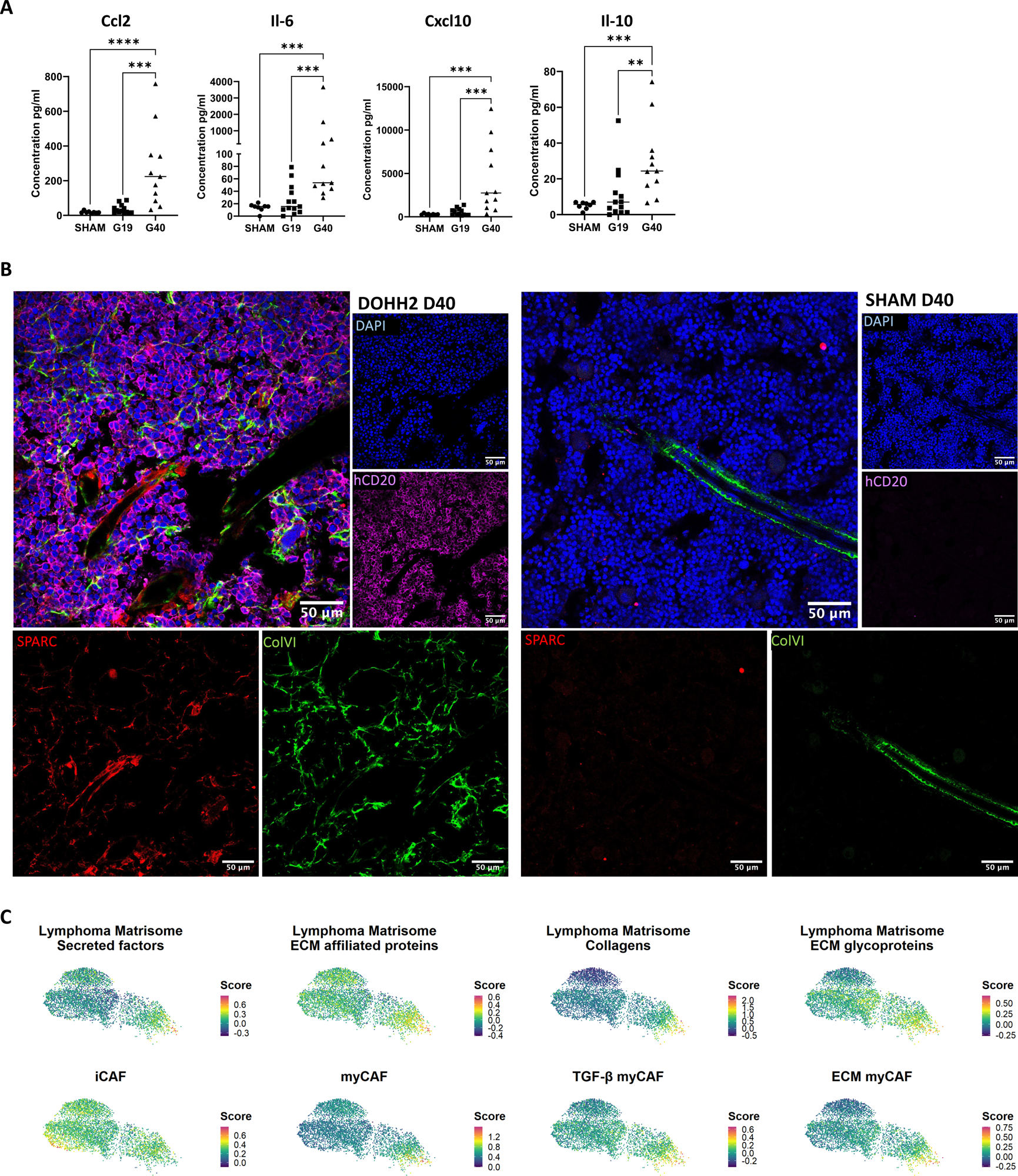
Validation of the stroma reprogramming in lymphoma-invaded BM. (A) Chemokine and cytokine levels were measured in BM plasma from Sham mice (n = 8) and DOHH2 grafted mice (D19 n = 13, and D40 n = 11) using MSD assay. ** P<.01, *** P<.001, and **** P<.0001. (B) BM sections of grafted (left) or Sham (right) mouse femurs were stained for hCD20 (magenta), Sparc (red), and Collagen VI (green). Nuclei were counterstained with DAPI (blue), Scale bars, 50 µm. (C) Lymphoma matrisome signatures previously defined from DLBCL LN (Kotlov *et al*., Cancer Discov 2021) and solid cancer CAF signatures previously defined from pancreatic cancer (Elyada *et al.*, Cancer Discov 2019) and from breast cancers (Kieffer *et al.*, Cancer Discov 2020) were plotted on the LepR^pos^ MSC clusters generated in Figure 3A.

Taken together, these results show the polarization of LepR^pos^ MSCs into an ectopic LSC-like and ECM remodeling phenotype upon direct contact with tumor B cells in mouse xenograft model, thus mimicking lymphoma CAF reprogramming.

### Study of the bidirectional B cell/stromal cell crosstalk in invaded BM

To explore how BM LepR^pos^ MSCs could progressively become more efficient to interact with and support lymphoma B cells and identify underlying mechanisms, we decided to decipher deeper the stromal cell differentiation program through a trajectory analysis based on the Slingshot algorithm^38^. We set the MSC_Ctrl as the starting point and MSC_Gr.Late as the end point for the trajectory generation, creating a pseudotime corresponding to the actual kinetic of the experiment, where MSCs from early grafted mice were found as an intermediate state between MSCs from control mice and MSC from late grafted mice (Figure 5A). The Tradeseq analysis of DEGs along pseudotime generated seven gene clusters (Figure 5B and Supplemental Table S3). Among them, clusters 4 and 1 included respectively 1242 and 2464 genes progressively upregulated over the pseudotime while cluster 2 included 1456 genes upregulated at the end of the pseudotime, collectively characterizing B-cell-primed MSCs. We then analyzed enrichment for transcription factor target genes in clusters 4, 1, and 2 using the TRRUST database (Figure 5C) and identified notably Smad3, Sox4, Srf, and Myod1^39–41^, all involved in the TGF-β signaling pathway, of as active at the end of the pseudotime (Figure 5D). We then performed a Nichenet interactome analysis using these B-cell primed MSCs as sender cells and DOHH2 at D40 as receivers (Figure 5E and Supplemental Figure S5A). Several members of the TGF-β family pathways were found enriched at the stroma cell/B-cell interface, including *Tgfβ1* and *Tgfβ3*, *Bmp2*, *Bmp3*, *Bmp5*, *Inhba*, and *Inhbb*, consistent with the progressive enrichment for the TGF-β myCAF signature in MSC_Gr.Early and MSC_Gr.Late (Supplemental Figure S4B). Moreover, adhesion molecules, ECM remodeling factors, and collagens were also found strongly enriched in this interactome study. Finally, the Notch ligand *Jag1* was also predicted as involved in the lymphoma BM niche. Interestingly, the 25 ligands that were the most strongly enriched at the interface between BM stromal cells and B cells essentially belonged to clusters 1 and 2, corresponding to genes expressed at the later time points of the pseudotime (Supplemental Figure S5B). Accordingly, similar data were obtained using MSC_Gr.Late as sender cells (Supplemental Figure S5C). Conversely, to better characterize the mechanisms of the lymphoma B cell-dependent reprogramming of BM LepR^pos^ MSCs, we applied the NicheNet computational method using DOHH2 D40 as sender cells and B-cell primed MSCs as receiver cells, focusing on ligand-receptor interactions (Figure 6A, left, and Supplemental Figure S6) and on ligand-targeted gene relationships (Figure 6A, right). In particular, *TGFB1* and *VEGF* expression by DOHH2 were predicted to contribute to ECM remodeling in BM stromal cells.

**Figure 5.**
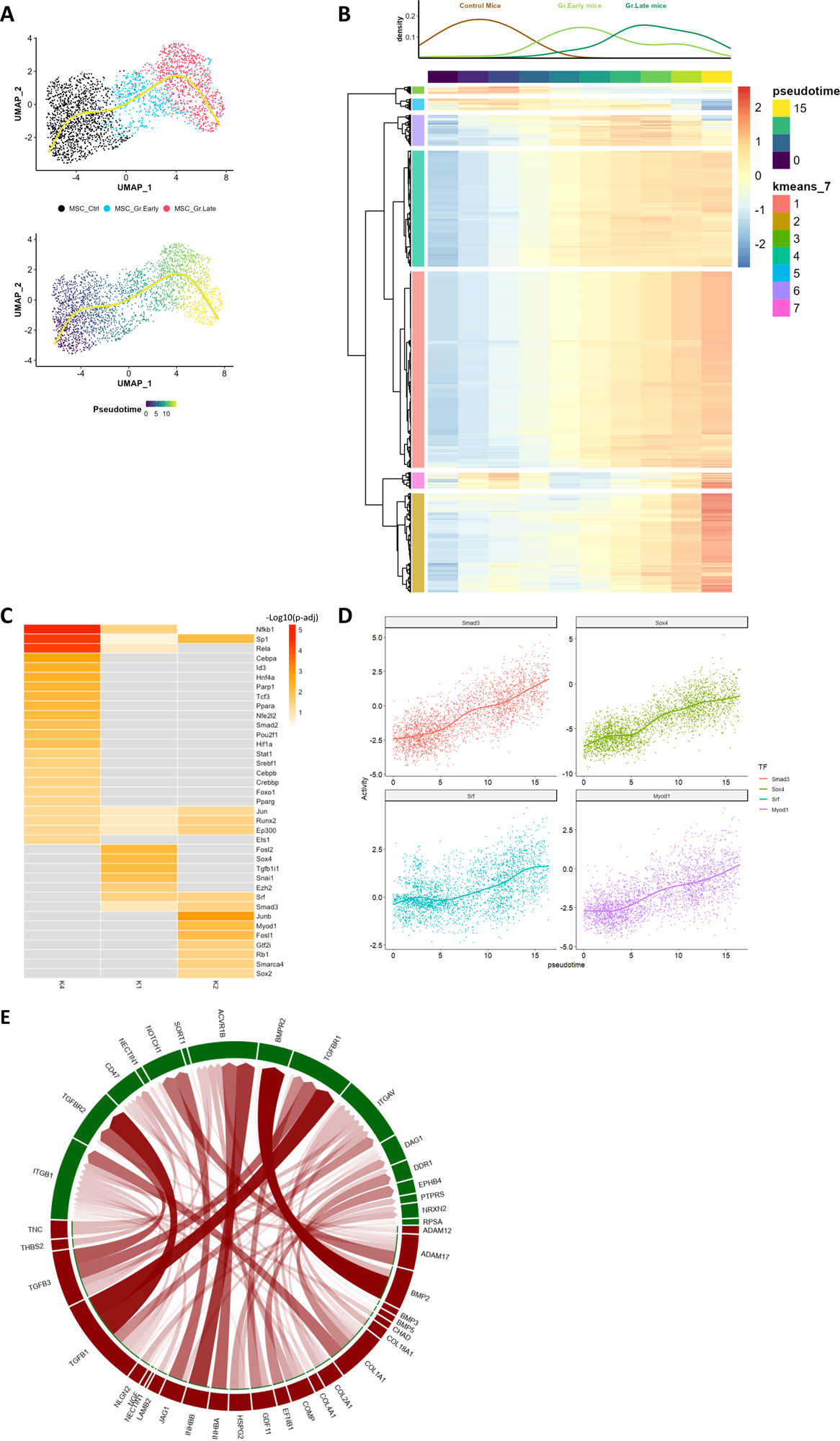
Impact of BM stromal cells on lymphoma B cells. (A) Pseudotime analysis of LepR^pos^ MSCs obtained by setting the MSC_Ctrl population as the starting point and the MSC_Gr.Late population as the end point. Yellow arrow represents the arbitrary obtained trajectory. In the lower panel, cells are colored according to their original DA-seq cluster. (B) Clustering of the 5,989 DEGs along the pseudotime highlighting 7 gene clusters. The upper panel shows the relative enrichment of control, Gr.Early, and Gr.Late experimental conditions along the pseudotime. (C) Transcription factors identified by TRRUST based on the enrichment of their targets among the TOP 200 differentially expressed genes in the clusters 1, 2, and 4 of the pseudotime were plotted on a heatmap. The color scale corresponds to the -Log10(p-adjust). (D) Activity of Smad3, Sox4, Srf, and Myod1 transcription factor along the pseudotime. Smoothed lines correspond to generalized additive model estimates. (E) Circos plots showing predicted interactions between LepR^pos^ MSCs and lymphoma B cells. B-cell-primed MSCs, corresponding to genes included in clusters 1, 2, and 4 upregulated over the pseudotime, were defined as senders and DOHH2 D40 as receivers.

**Figure 6.**
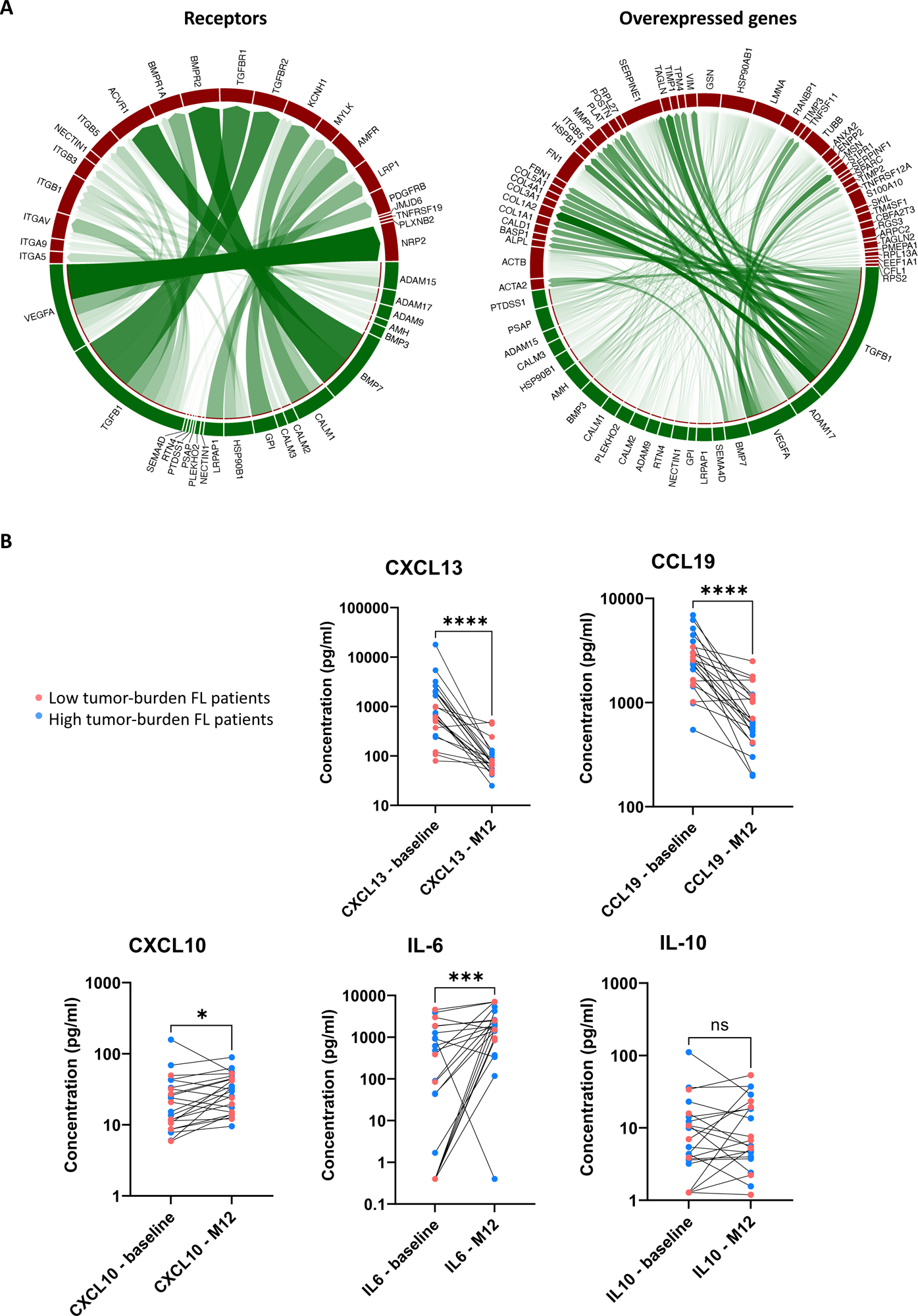
Impact of lymphoma B cells on BM stromal cells. (A) Circos plots showing predicted interactions between DOHH2 cells and LepR^pos^ MSC at the late stage of graft. DOHH2 recovered from the grafted mouse at the late time point (DOHH2 D40) were defined as senders and B-cell-primed MSCs, corresponding to genes included in clusters 1, 2, and 4 upregulated over the pseudotime as receivers. (Left Panel) Interaction between ligands from DOHH2 D40 and predicted receptors from B-cell-primed MSCs. (Right Panel) Interaction between ligands from DOHH2 D40 and ligands-derived upregulated target genes (downstream signals induced by ligands-receptors crosstalk) from B-cell-primed MSCs. (B) Quantification by Luminex of cytokine and chemokine levels in BM plasma from FL patients (n=22) collected at diagnosis and after 12 months of treatment. Blue dots represent high tumor-burden FL patients and pink dots represent low tumor-burden FL patients * P<.05, *** P<.001, and **** P<.0001.

We then wondered whether the reprogramming of BM stromal cells by tumor B cells is reversible upon lymphoma cell clearance. To address this issue, we used BM plasma collected from BM-invaded FL patients at diagnosis and one year after treatment with a strong reduction of BM infiltration (Figure 6B). Upon treatment, levels of LSC chemokines CXCL13 and CCL19 were significantly decreased, including in low-tumor burden patients treated by Rituximab only (FLIRT trial, pink dots), *i.e*. with a therapeutic activity restricted to B cells. IL-10 levels remained at the same level even after treatment. Unexpectedly, IL-6 and CXCL10 inflammatory mediator levels increased in BM plasma after treatment, suggesting a partial maintenance of the deregulated BM niche. Altogether, these data shed new lights on the molecular pathways underlying stromal cell/B cell crosstalk with invaded BM.

## Discussion

With this work, we provided additional clues on the induction of an ectopic local LSC population with tumor supportive properties upon FL B-cell BM invasion. Indeed, by looking at the chemokine levels within FL patient BM plasma, we observed an upregulation of the LSC specific chemokines CCL19 and CXCL13, compared with HD. These chemokines are classically produced by LN LSC, respectively FRCs and FDCs. This phenotype was related to the LSC-inducer cytokine TNF, and to the FL-CAF inducer TGF-β, that increased within BM plasma of FL patients as recently reported within FL LN where they support the crosstalk between FL B cells and stromal cells and synergize to increase the transcriptional level of CCL19/CCL21^17^. Of note, FL-derived EVs are also able to prime BM stromal cells without direct cell interaction^4^ and this effect is also dependent on TGFβ signaling, reinforcing the role of this factor in the biology of B-cell lymphomas besides its previously proposed effect on lymphoma-infiltrating cytotoxic T-cell immunosuppression^42,43^. We also provided a first characterization of the ECM remodeling within FL-invaded BM, with increased Collagen VI networks within FL B-cell invaded areas and identified, using our xenograft model, a direct effect of tumor B-cell invasion in this process. Interestingly, not only BM stromal cells, but also DOHH2 themselves overexpressed ECM-related factors after *in situ* xenograft and could contribute to the local ECM modifications. This is reminiscent of the recent demonstration that culture of lymphoma B-cell lines in 3D spheroids increases their production of ECM components^29^, something that was neglected in classical 2D *in vitro* experiments. Interestingly, DOHH2 cells at D40 also upregulated *ITGA4* and *ITGB1*, together forming the VLA-4 integrin that upon ligation with VCAM-1 (expressed in MSC_Gr.Late) was associated with GC B-cell lymphoma growth and resistance to the anti-CD20 antibody Rituximab^44^. Of course, non-malignant immune compartments are missing or dysfunctional in our immunocompromised mouse models and could play a crucial role in lymphoma ECM reorganization. In particular in DLBCL LN, macrophage-derived ECM was shown to be strongly modified and contribute directly and indirectly to tumor cell growth^35^. The upregulation of CCL2 in both xenografts and FL patients could contribute to the recruitment and further polarization of such tumor-infiltrating ECM-producing macrophages^18^. Conversely, ECM network alterations could impact anti-lymphoma immune cell response^25^. Humanized mouse models allowing the development of fully functional lymphoid structures are urgently needed to further explore this major question.

Whereas the phenotype of lymphoma-primed BM stromal cells was strongly different from that of normal B-cell primed BM stromal cells previously described in steady-state immunocompetent mice, it was at least partly share in various mature B-cell hematological malignancies, including MM. Moreover, similarity in ECM remodeling was found between lymphoma-primed BM stromal cells and DLBCL LN. Such ECM remodeling was also explored in a subcutaneous xenograft mouse model of DLBCL in immunocompromised mice^26^. In this study, deregulation of Jun pathway in tumor B cells was found to be involved in the expression of ECM adhesion (such as integrins), surface receptors (CCR5, CCR7, IL-6R), ECM modeling enzymes (metalloproteinase) and cytokines (IL-10, IL-6, IL-7) supporting the capacity of DLBCL cells to interact with ECM and stromal cells and to home within BM^26^. In accordance, Jun deficient tumor cell lines exhibited a lower tumor volume and a decreased BM infiltration^26^. More recently, B-cell derived TNF and LT (previously described as non-redundant LSC inducers^20^) has been involved in FRC elongation in DLBCL, showing another conserved stromal cell polarization effect by malignant FL and DLBCL B cells^25^. Hence, this suggests that the effect of malignant B cell on stromal cell polarization is largely conserved, affecting the ECM remodeling and the secretory profile of the MSCs. Consistently, the secretory profile might also be conserved in polarized BM stromal cells among mature B-cell derived malignancies. Indeed, MM MSCs, like our FL BM stromal cells overexpress *CCL2, IL-6*, *COL6A3,* and *FN1*^34^. Again, our model does not recapitulate some aspects of cytokine/chemokine deregulation in lymphoma BM. In particular, we previously demonstrated that IL-4, mainly produced by CD4^pos^ T cells, is involved in the upregulation of CXCL12 within FL LN and FL BM whereas the contact with tumor B cells reduces CXCL12 expression in vitro in a TNF-dependent manner^21^. No model is currently available to evaluate such complex niche in vivo.

The remodeling of the ECM might exert several protumoral effects, favoring the implantation of FL B cells within certain areas of the BM and promoting the release of chemokines and cytokines. Interestingly, it has been demonstrated that heparanase, an endoglycosidase triggering ECM disassembly may increase the bioavailability of growth factors bound to heparin sulfates including CXCL12, and is strongly expressed in about 50% of FL and DLBCL^45^. Moreover, clues might indicate that this heparinase activity is due to the local microenvironment rather than malignant B cells^45^. Furthermore, many growth factors and cytokines/chemokines are trapped within the ECM such as TGF-β, Fibroblast growth factor (FGF) or CXCL12 ^45,46^ and might be released upon ECM remodeling. CCL2, CCL5 or CXCL9 and CXCL10 might also be released upon ECM degradation and reshape^47^. CCL2 have already been discussed as associated with TAM recruitment in FL^18^ while CCL5 in breast cancer has been associated to enhanced metastasis and tumor dissemination^48^, and is secreted by MSCs^49^. Finally, integrin interaction with ECM molecules can trigger “outside-in signaling”, that may activate several intracellular pathways such as the actin cytoskeleton or the cell cycle^50^. Such interactions influence cell migration (and homing) and survival^50^. Moreover, ECM reshape within the FL BM might also affect effector immune cells invasion and tumor elimination.

An important open question is whether this polarization of BM stromal cells into FL-CAF is reversible or not. By looking to our last results, tumor B-cell decrease within BM is associated with a downregulation of the LSC-associated chemokines CCL19 and CXCL13, thus suggesting a reversion of the LSC phenotype. However, CXCL10 levels are slightly increased after the treatment showing that phenotype reversion might be incomplete or that a different phenotype is induced after tumor elimination. This CXCL10 increase might be helpful to recruit NK cells or CD8+ T cells to eliminate the possibly remaining tumor cells, or Tregs to favor immune suppression and tissue repair^51,52^. Thus, it is possible that part of the BM supportive stroma remains as a residual niche supporting treatment-resistant lymphoma B cells. These stromal cells might thus facilitate FL B cell reseeding as FL is still incurable and characterized by multiple relapses^3,53^. Deeper characterization of this niche is hence required to better understand how FL relapses, to prevent such events and ameliorate patient prognosis.

## Materials and Methods

### Human samples and cell lines

The research protocol was conducted under French legal guidelines with informed consent and was approved by the local ethics committee. BM aspirates were obtained from low-tumor burden FL patients (FLIRT trial, NCT02303119), and high-tumor burden FL patients (GATA trial, NCT03276468 and EpiRCHOP trial, NCT02889523), all collected at diagnosis and, for a part of them, after 12 months of treatment. BM plasma was collected by centrifugation and frozen at -80°C until use. BM aspirates were also collected from age-matched patients undergoing cardiac surgery and used as HD. BM biopsies were obtained from patient diagnosis and reviewed by expert pathologist as FL or normal BM. Specimens were fixed in formol, rinsed and placed in decalcifier working solution. Following decalcification, the BM biopsies were processed and embedded in paraffin wax. Four-μm thick sections were cut using a microtome for further use in immunohistofluorescence studies.

The FL cell line DOHH2 was obtained from the DSMZ cell collection (Braunschweig, Germany) and the GCB-DLBCL cell line OCI-Ly-19 was a gift from Dr Lou Staudt (National Cancer Institute, Bethesda, MD). DOHH2 cells were maintained in RPMI (Gibco) with 10% fetal bovine serum (FBS, Eurobio) and OCI-Ly-19 cells in IMDM with heat inactivated 10% human AB serum (Biowest), 50 µM 2-Mercaptoethanol (Sigma Aldrich) and 1x penicillin/streptomycin. Luciferase-expressing DOHH2 cells (Luc-DOHH2) were established after transduction with a lentivirus carrying the pHAGE PGK-GFP-IRES-LUC-W plasmid (gift from Darrell Kotton, Addgene plasmid # 46793; http://n2t.net/addgene:46793; RRID:Addgene_46793), which contained the enhanced GFP and luciferase genes under the control of PGK promoter^54^. Cells were transduced in RPMI supplemented with 10% FBS and 8 μg/ml protamine sulfate (P4020, Sigma Aldrich). Virus was added at a MOI 5 and a spinoculation step (centrifugation x1000g at 32°C for 30 min) was performed to enhance lentiviral transduction. After 24 hours, the medium was replaced with standard culture medium. After 1 week, Luc-DOHH2 cells were isolated using a cell sorter based on their co-expression of GFP (FACS Aria II, BD Biosciences).

### Xenograft models

Mice were maintained in specific pathogen-free conditions in Rennes animal facility. All procedures were carried out with the approval of the Committee on the Ethics of Animal Experiments under the French Ministry of Higher Education and Research (permission#: 8556– 2017011613335049_v8) and in strict accordance with the recommendations in the Guide for the Care and Use of Laboratory Animals, EEC Council Directive 2010/63/EU. Rag2^-/-^γ_c_^-/-^ mice on a C57BL/6 background were a gift from James Di Santo (Pasteur Institute, Paris, France)^55^. Seven to ten-week-old male Rag2^-/-^γ_c_^-/-^ mice were transplanted with 0.5 x 10^6^ DOHH2, OCI-Ly-19, or Luc-DOHH2 cells by intrafemoral route or were injected with PBS (Sham mice). Two days before intrafemoral grafts, mice underwent a myeloablation using two intraperitoneal injections of hydroxyurea (H8627, Sigma Aldrich, 1 mg/g) separated by 8h. For intrafemoral injection, mice were anesthetized by intraperitoneal injection of a mix of 75 mg/kg ketamine and 8 mg/kg xylazine. A hole was done using a G25 needle and 30 µl of cell suspension in PBS or PBS alone was slowly injected into the intrafemoral space using a G26 needle with a 1 ml syringe. Mice were then monitored every day until sacrifice. DOHH2 infiltration in blood, spleen, and femurs (injected and contralateral) was studied by flow cytometry on a CytoFLEX (Beckman-Coulter™) at different timepoints as the percentage of hCD20^pos^ cells (anti-human CD20 FITC, clone 2H7) compared to murine CD45^pos^ cells (anti-mouse CD45 PE, clone 30F11, ThermoFisher). When indicated, tumor dissemination of Luc-DOHH2 was monitored using a PhotonImager Optima system (Biospace Lab) equipped with a camera box and a warming stage. Monitoring of the *in-situ* bioluminescence was performed by 100 mg/kg intraperitoneal injection of D-luciferin (FP-M1224D, Interchim) and mice imaging with an exposure time of 1 min. To collect BM plasma, the proximal epiphysis of the injected femur was cut and removed. Femur was placed in a 0.5 ml microcentrifuge tube bearing a small hole at the bottom and nested in a heparinized 1.5 ml microtube. After a centrifugation step (6,500 × g for 1 min) BM plasma was recovered into a new collection tube and frozen at -80°C until analysis. Injected BM femurs were also collected for immunofluorescence imaging.

### Immunofluorescence imaging

Mouse femurs were fixed during 24h in 4% paraformaldehyde, incubated sequentially in 10%, 20% and 30% sucrose (1h each), embedded in SCEM embedding medium (Section-Lab) and frozen at -80°C. Then, 18 μm cryosections of undecalcified femoral bone were generated using Kawamoto tape transfer method^56^. For immunofluorescent staining, sections were rinsed with PBS and blocked with 2% donkey serum and 1% bovine serum albumin in PBS containing 0.1% Triton X-100 and 0.005% Tween20 for 1h at RT. Slides were then incubated in a humidified chamber overnight at 4°C with primary antibodies (1:50 dilution): rabbit anti-human CD20 (ab78237, Abcam), rat anti-mouse Fibroblast Marker ER-TR7 (sc73355, Santa Cruz). After washes, slides were incubated for 1h at RT with secondary antibodies used at 1:250 dilution: anti-rat 405 (Jackson Lab), anti-rabbit 594 (Jackson Lab), anti-mouse 488 (Jackson Lab). Finally, tissue sections were mounted with Mowiol (Merck) antifade reagent containing DAPI (Sigma Aldrich) and analyzed by confocal microscopy on a SP8 (Leica Microsystems). ImageJ software (National Institutes of Health) was used for image analysis.

Immunofluorescence on human BM biopsies were performed on VENTANA Discovery automated staining instrument according to manufacturer’s instructions. After deparaffinization and epitope retrieval, slides were incubated with primary antibodies: rabbit anti-PAX5 AF647 (EPR3730, Abcam); rabbit anti-Collagen VI AF594 (EPR17072 Abcam) and mouse anti-SPARC/Osteonectin (ON1-1, Takara) for 1 h at RT. Slides were then washed 4X with TBS containing 0.03% Triton-X. Then, the slides were incubated with anti-mouse-AF488 for 45 min at room temperature. The slides were then washed 4X with TBS containing 0.03% Triton-X. One drop of VECTASHIELD Antifade Mounting Media (Vector Labs, Burlingame, CA) containing DAPI (4’,6-diamidino-2-phenylindole) was then added to the slides prior to the placement of the coverslip.

### Cytokine and chemokine quantification

Human cytokine and chemokine levels in BM plasma were measured by Luminex technology. The following proteins were analyzed: TGF-β (FCSTM17, R&D Systems), CCL2 and TNF (HCYTOMAG, Merck Millipore), CCL19, CXCL10, CXCL13, IL-6, and IL-10 (ProcartaPlex, Thermo Fisher) according to manufacturer’s instructions. For TGF-β quantification, samples were diluted at 1/15. Analysis of the mouse cytokines and chemokines was carried out using a Meso Scale Discovery (MSD) mouse U-PLEX Custom biomarker group 1 and was realized by the InnovaBIO platform (CHU Caen, France). Values under the detection limits established by the manufacturers were replaced by their detection limits for each parameter.

Statistical tests and graphic representation of cytokine and chemokine quantification results were performed with Prism software version 9.5.1 (GraphPad Software). Statistical significance was determined using the Mann-Whitney nonparametric U test for Human samples. Correlation between CXCL13 and CCL19 concentrations was performed using a Spearman test. For mice samples, statistical significance was determined using Kruskal-Wallis non-parametric test with uncorrected Dunn’s multiple comparisons to compare each condition side by side with the others.

### Purification of murine endothelial and stromal BM cells and DOHH2 cells

Mouse femurs of steady-state (control), PBS-injected (Sham), and DOHH2-injected (grafted) mice were harvested at D19 and D40. BM was flushed and dissociated in PBS containing DNase I (10 U/ml, Pulmozyme®, Roche) and Liberase DH (0.025 mg/ml, 5401089001, Roche). Bones were cut into small fragments using scissors in a dissociating media containing 10 U/ml DNase I and Collagenase P (1.2 mg/ml, 11213865001, Roche). Bones and BM were digested at 37°C for 20 min under agitation. PBS supplemented with 5% FBS and 15% sodium citrate was added to neutralize the digestion solution, followed by a centrifugation at 450 g for 5 min. After filtration on 100 µM cell strainer, depletion of red blood cells was carried out with RBC lysis buffer (00-4333-57, Thermo Fisher), stromal cell enrichment was performed using anti-Cd45 (clone 30-F11, 103102, Biolegend) and anti-Ter119 (116202, Biolegend) Abs and using magnetic beads (Dynabeads) according to manufacturer’s instructions. After depletion, cells were stained for sorting with the following antibodies: anti-Cd31 BUV395 (clone MEC13.3, BD Biosciences), anti-Cd51 BV650 (clone RMV-7, BD Biosciences), anti-Sca1 PECy7 (clone D7, Biolegend), anti-Cd200 APC (clone OX-90, Biolegend), anti-Cd45 APC-Cy7 (clone 30F11, BD Biosciences), anti-Ter119 APC-Cy7 (116223, Biolegend), anti-hCD45 APC-Cy7 (clone 2D1, BD Biosciences) and anti-hCD20 FITC (clone 2H7, 555622, BD Biosciences) and with live dead reagent (L34959, Thermo). Cells were then sorted using a FACSAria II (BD Biosciences) as Cd45/Ter119^neg^Cd31^neg^Cd51^pos^Cd200^pos^ stromal cells, Cd45/Ter119^neg^Cd31^pos^Sca1^pos^ endothelial cells^57,58^, and hCD20^pos^ human lymphoma B cells.

### DOHH2 RNA sequencing and analysis

After cell sorting, BM DOHH2 were resuspended in RA1 buffer and tris(2-carboxyethyl)phosphine (TCEP) from Macherey-Nagel and stored at -80°C for less than 3 months. RNA was extracted using NucleoSpin RNA kit (Macherey-Nagel) and RNA integrity number (RIN) was assessed using an RNA 6000 Nano kit (5067-1511, Agilent) on a 2100 Bioanalyzer from Agilent. All RIN were above 7. RNA was sequenced on a NovaSeq 6000 (Illumina). Raw sequencing read data was checked and visualized for quality using FastQC (v0.11.5). Reads were aligned to the reference genome Hg38_R90 using STAR (v2.5.2a). Read summarization, gene and transcript counts were performed using the featureCounts (v1.4.6). Analysis of differentially expressed genes (DEGs) was performed using DESeq2 (v1.40.1) R package^59^. P values were adjusted with FDR Benjamini-Hochberg procedure. Log2 fold changes (Log2FC) were moderated using Lfcshrink method. Genes were ranked according to the Log2FC, and we performed a pre-ranked Gene Set Enrichment analysis (GSEA^60^) analysis to explore MSigDB Hallmarks, Canonical pathways, and Gene Ontology databases. GSEA pathways with an FDR < .05 were plotted when relevant to lymphoma biology. The results were plotted using R studio and ggplot2 package.

### Single-cell RNA sequencing and analysis

Sorted stromal and endothelial cells were merged at a 1/1 ratio before encapsulation into emulsion droplets using Chromium Controller (10x Genomics) with a target output of ∼20,000 cells. scRNAseq libraries were constructed using Chromium Single Cell 3’ v3.1 Reagent Kit according to the manufacturer’s recommendations. Library quality control and quantification were performed using a KAPA Library Quantification kit for Illumina platforms (Kapa Biosystems, KK4873) and a 2100 Bioanalyzer High Sensitivity DNA kit (Agilent, 5067-4626). Libraries were sequenced on an Illumina HiSeq X Ten system with an average depth of 31,439 reads per cell, then mapped to the mouse genome (build GRCm38) and demultiplexed using CellRanger pipelines (10X Genomics, v.3.1.0).

Reads were aligned to the mouse genome and converted to count tables using Cell Ranger count (v5.0.0; 10X Genomics). The resulting expression matrices were processed individually in R using Seurat package^61^ (v4.3.0). First, cells were filtered according to classical QC metrics (number of UMI and detected genes per cell, proportion of mitochondrial reads) using specific cutoffs. In particular we excluded cells with fewer than 1,000 detected genes and more than 8,000 detected genes and with more than 10% mitochondrial reads. All the samples were then merged and processed using the standard Seurat workflow (NormalizeData, ScaleData, FindVariableFeatures, RunPCA, RunUMAP, FindNeighbors and FindClusters functions). DEGs between clusters or conditions were inferred with Wilcoxon Rank Sum test as implemented in the FindAllMarkers function. Enrichment of experimental conditions in scRNA-seq clusters was performed with a Fisher’s exact test with the alternative hypothesis set to “greater”.

Processed scRNA-seq datasets from C57BL/6 mouse BM non-hematopoietic cells were downloaded from the Broad Institute single cell portal (https://portals.broadinstitute.org/single_cell/study/mouse-bone-marrow-stroma-in-homeostasis)^24^. DEGs in each cluster were downloaded from the original article and filtered (based on average log fold change) to generate Top-50 gene signature scoring. Signature scores were computed with the AddModuleScore function of the Seurat R package. This function calculates for each individual cell the average expression of each gene signature, subtracted by the aggregated expression of control gene sets. Enrichment of clusters for specific gene signatures was performed using Student t-test with p-value adjustment by Holms correction.

Cell clusters with differential abundance (DA) between one condition against the others were computed using the DA-seq v1.0.0 package^31^. Briefly, DA-seq predicts DA scores for each cell under two separate conditions by applying a logistic regression model. Label permutation is then used to empirically evaluate the statistical significance of the prediction results. Each DA-seq cluster was filtered to exclude outlier cells not belonging to the corresponding unsupervised Seurat cluster (Supplemental Figure S7).

### Trajectory analysis

Trajectory inference was performed using the slingshot (v2.5.2) package^38^ on MSC_ctrl, MSC_Gr.Early, and MSC_Gr.Late DA clusters. Slingshot was run on a UMAP embedding these three clusters and the MSC_ctrl DA cluster was manually designated as the root of the trajectory. Trajectory-based differential expression analysis was performed with the tradeSeq (v1.7.07) package^62^. Gene expression was smoothed along the trajectory with the fitGAM function and the associationTest function was then applied to detect DEGs over the pseudotime.

### Interactome analysis

For CellPhoneDB analysis^63^, stromal cell clusters were selected from our dataset and from already published murine BM stromal cell dataset^24^. Gene names were converted to their human equivalent with the convert_mouse_to_human_symbols function from the Nichenetr package^64^. Raw scRNA-seq data from human FL B cells^33^ were downloaded from European Genome-Phenome Archive (accession number EGAS00001006052) and processed with Cell Ranger count (v7.0.0; 10X Genomics). Resulting count tables were filtered and annotated in accordance with the processed dataset available in the Chan Zuckerburg cellxgene repository. Tumor B cell clusters were filtered out and merged with stromal cell clusters. Corresponding count table was normalized according to author recommendations and statistical inference of interaction specificity was performed (CellPhoneDB v4 method 2).

Interactome analysis between DOHH2 cells and BM-MSCs was performed using the Nichenetr (v1.1.1) package. First, we used the genelist obtained by combining clusters 1, 2, and 4 from the pseudotime analysis, *i.e.* genes that were induced along the pseudotime, to generate B-cell-primed MSCs as sender cells. Interactions were then predicted between sender (B-cell-primed MSCs or MSC_Gr.Late cells) and receiver cells (DOHH2 D40) to explain the upregulated DEGs between DOHH2 D40 and DOHH2 D0. Conversely, we also defined DOHH2 D40 (based on the DEGs compared to D0) as senders and B-cell-primed MSCs as receivers. For DOHH2, DEGs were obtained from RNAseq data by comparing D40 versus D0 with an FDR of 5% and a log2 fold change > 0.58. Expressed genes were defined based on a mean average log2 expression above 6 in D40 condition. For MSC_Gr.Late cells, differentially expressed genes were obtained by comparing this cluster with the MSC_Sh clusters with an FDR of 5% and a log2 fold change > 0.5. In addition, we used a bootstrap test with 1000 random gene permutations (among the differentially expressed genes in receiver cells) to create a distribution of each ligand activity and estimate a P-value for each NicheNet observed activity.

### Inference of transcription factor activity

Key transcriptional regulatory networks supporting MSC pseudotime were identified using TRRUST database (https://www.grnpedia.org/trrust/) using the Top 200 upregulated genes (P-adj < .05) in clusters 1, 2, and 4 of the pseudotime. The log10(p-adjust) values were then visualized through an heatmap using pheatmap package (v1.0.12). Inference of transcription factor activity was then performed using the dorothea package (v1.10.0; https://saezlab.github.io/dorothea/). The mgcv package^65^ was used to smooth TF activities along the pseudotime using generalized additive models.

### scRNAseq Graphical representation

scRNA-seq data visualizations was produced with the Seurat, ggplot2, pheatmap, patchwork and ktplots packages.

Ligand-target interactions were represented through a circos visualization (circlize^66^ R package, v0.4.9). Ligands and receptors were then color-tagged according to their cell of origin. Links between ligands and targets were drawn with a transparency scale proportional to ligand-receptors interaction weights.

### Data availability

scRNAseq data are available on GEO ncbi database.

## Acknowledgements

This work was supported by research grants from the Institut National du cancer (INCA AAP PNP19-009), and Ligue Nationale contre le Cancer (Equipe Labellisée). BB is supported by the Rennes University and the Région Bretagne. High-throughput sequencing has been performed by the ICGex NGS platform of the Institut Curie supported by the grants ANR-10-EQPX-03 (Equipex) and ANR-10-INBS-09-08 (France Génomique Consortium) from the Agence Nationale de la Recherche (“Investissements d’Avenir” program), by the ITMO-CANCER and by the SiRIC-Curie program - SiRIC Grant « INCa-DGOS-4654 ». Some bone marrow plasma were obtained from patients included in the BIO-FLIRT study as a part of the ancillary study of the FLIRT trial (NCT02303119), in the BIO-GATA study as a part of the ancillary study of the GATA trial (NCT03276468), and in the BIO-EpiRCHOP study as a part of the ancillary study of the EpiRCHOP trial (NCT02889523), all sponsored by the Lymphoma Academic Research Organisation (LYSA). MSD quantification of murine soluble factors in BM plasma was performed by the InnovaBIO platform (CHU Caen, France). The authors thank the staff of the flow cytometry, animal housing, MRic Photonics, and Histo-Pathology Hight Precision core facilities (UMS Biosit, Rennes, France)

## Author contributions

ED: designed and performed experiments, analyzed data, and contributed writing; BB: performed experiments, analyzed data, and wrote the paper; SL: coordinated and performed bioinformatic analyzes; CM, TL, JD, FJ: performed experiments; F L-G: provided FL samples; DR, FM: analyzed data and discussed results; KT: designed and supervised research, analyzed data, and wrote the paper.

