## Supplemental Figures for "Lymphoma B cells remodel bone marrow stromal cell organization and function to induce a supportive cancer-associated fibroblast network"

#### Supplemental Figure S1.

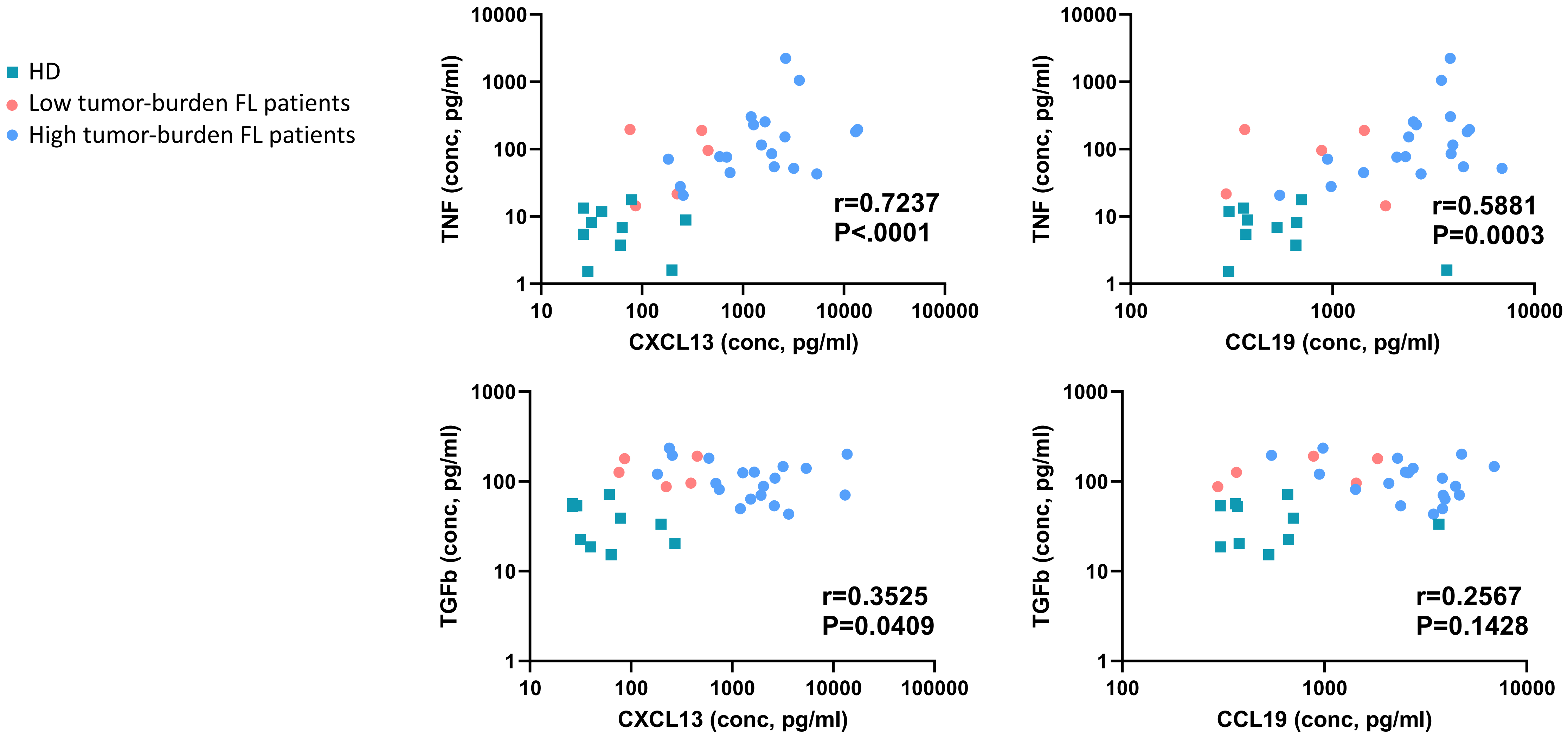


**Supplemental Figure S1. Correlation of soluble factors in BM plasma (related to Figure 1)**

Correlations between TNF or TGFβ1 level and CXCL13 and CCL19 levels in BM plasma from FL patients (n=24) and HD (n=10) determined by Luminex assay. Green squares represent HD, blue dots represent high tumor-burden FL patients, and pink dots represent low tumor-burden FL patients.

#### Supplemental Figure S2.

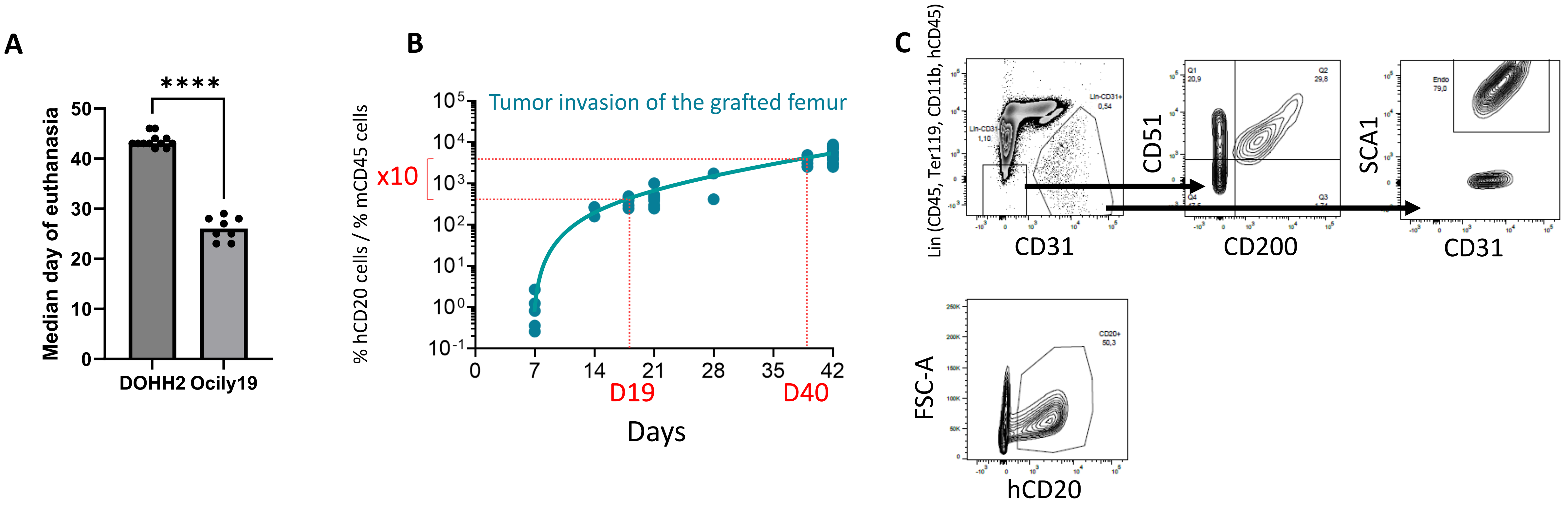


**Supplemental Figure S2. Characterization of DOHH2 xenograft model (related to Figure 2)**

(A) Median day of euthanasia for RAG$^{-/-}$γc$^{-/-}$ mice after intrafemoral injection of 0.5x10$^{6}$ DOHH2 or OCI-Ly-19 cells. **** P< .0001

(B) Evolution over time of the proportion of viable human tumor cells (hCD20$^{pos}$) in the grafted femur compared to murine CD45$^{pos}$ cells. The early time point, day 19 (D19), corresponded to 10-fold less tumor invasion than the late time point, day 40 (D40).

(C) Gating strategy for the sorting of BM stromal and endothelial cells at D19 and D40 post-graft (upper panels). Gating strategy for the sorting of DOHH2 tumor cells at D19 and D40 post-graft (lower panel).

**Supplemental Figure S3.**

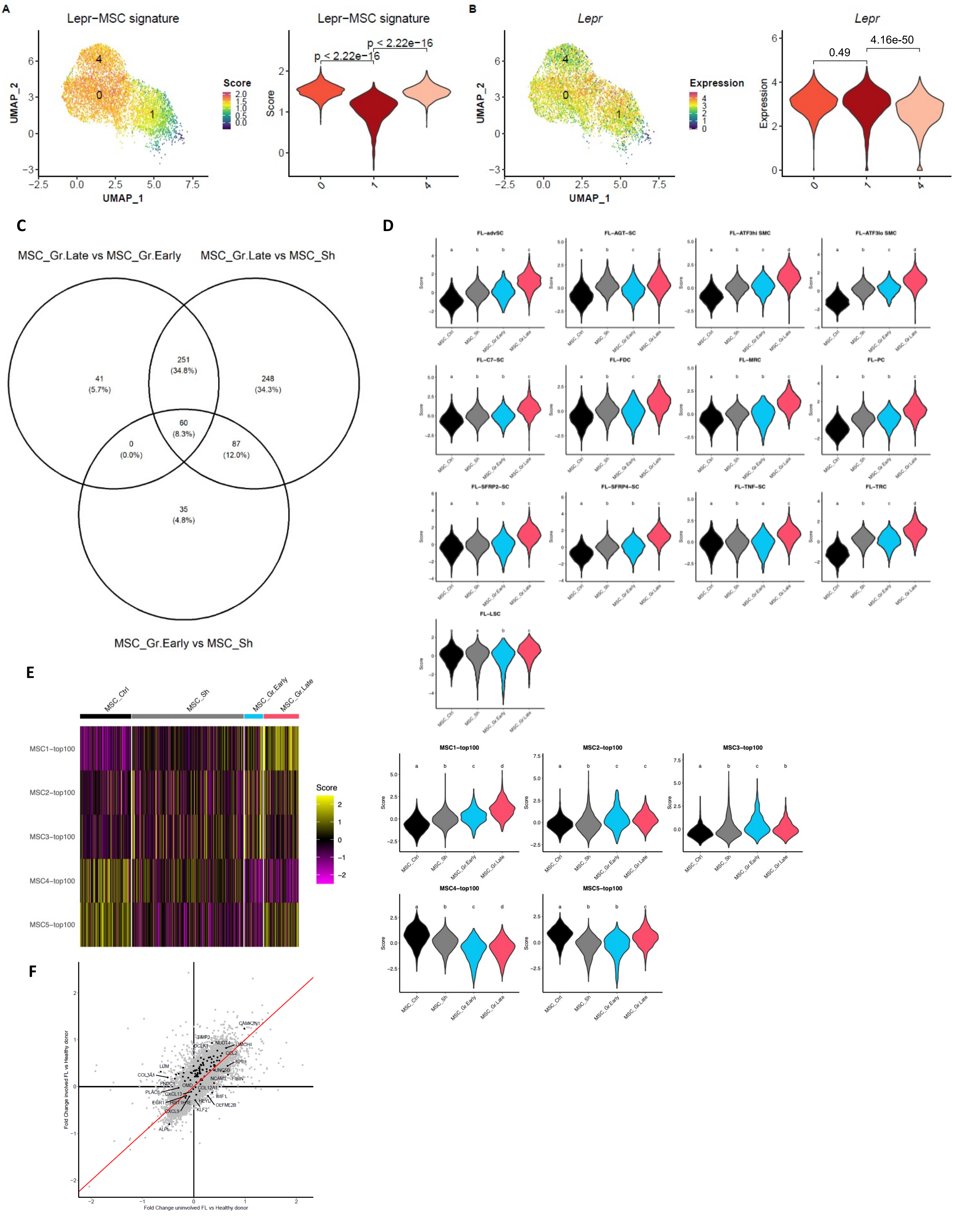

**Supplemental Figure S3. Lymphoma B cells drive LepR$^{pos}$ MSC transcriptional reprogramming (related to Figure 3).**

(A) Lepr-MSC signature previously defined by scRNAseq from steady-state C57BL/6 mice (Baryawno *et al.*, Cell 2019) was plotted on the LepR$^{pos}$ MSC clusters generated in Figure 3A. Corresponding Seurat scores were visualized on violin plots. P-values were defined with a Student t-test.

(B) Expression of *LepR* on the LepR$^{pos}$ MSC clusters generated in Figure 3A. Seurat scores were visualized on violin plots. Adjusted P-values were computed by the FindMarkers function from the Seurat package with default parameters.

(C) Venn diagram of the DEG (padj <.05; absolute value of Log2FC>0.25) between MSC_Gr.Late vs MSC_Gr.Early, MSC_Gr.Late vs MSC_Sh, and MSC_Gr.Early vs MSC_Sh.

(D) Specific gene signatures of LN FL-LSC subsets previously defined by scRNASeq (Abe et al., Nat Cell Biol 2022) were compared to our 4 DA-seq generated MSC clusters generated in Figure 3C. Corresponding Seurat scores were visualized on violin plots. Letters indicate statistically different conditions as defined by Student t-test with p value adjustment by Holms correction.

(E) Heatmap of gene expression values from the BM MSC clusters obtained from HD and MM patients (published in de Jong *et al.* Nat Immunol 2021), relative to the LepR$^{pos}$ MSC clusters identified by DA-seq. Corresponding Seurat scores were visualized on violin plots. Letters indicate statistically different conditions as defined by Student t-test with p value adjustment by Holms correction.

(F) Scatterplot of genes significantly modulated in BM-MSCs obtained from FL patients with a BM involvement (FLE, y axis) or not (FLN, x axis) compared to BM-MSCs obtained from HD highlighting the 100-genelist described in Figure 3G. Gene expression profile of a previous cohort of FL BM-MSCs (Guilloton *et al.*, Blood 2012) was reanalyzed from normalized expression data (provided by the original authors) with the idep website (Xjin Ge et al., BMC Bioinformatics 2018).

### Supplemental Figure S4.

A

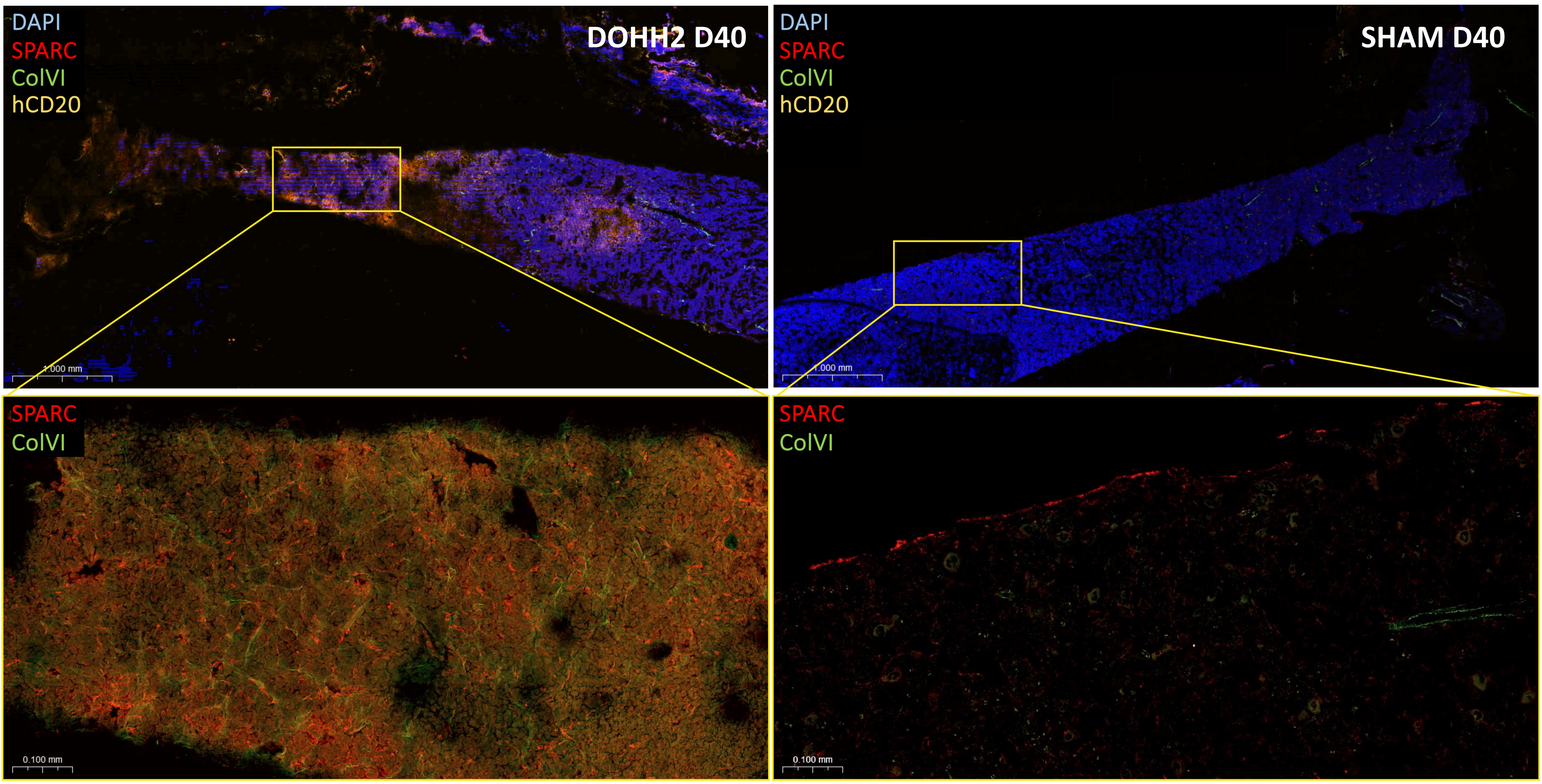


B

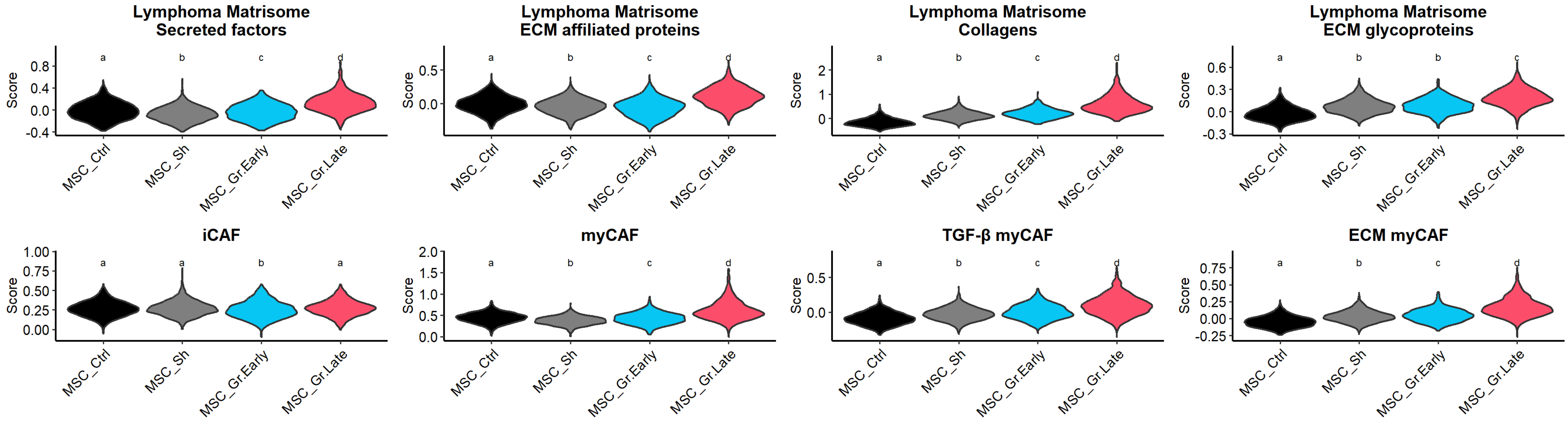


**Supplemental Figure S4. Validation of ECM deregulation in lymphoma-invaded BM stromal cells (related to Figure 4).**

(A) BM sections of grafted (Up) or Sham (Down) mouse femurs were stained for hCD20 (yellow), SPARC (red), and Collagen VI (green). Nuclei were counterstained with DAPI (blue). Scale bar, 1 mm. Boxes indicate areas magnified in the bottom panels where scale bars represent 0.1 mm.

(B) Lymphoma matrisome gene signatures previously defined from DLBCL LN (Kotlov *et al.*, Cancer Discov 2021) and CAF gene signatures previously defined from pancreatic (Elyada *et al.*, Cancer Discov 2019) and breast (Kieffer *et al.*, Cancer Discov 2020) cancers were compared to the four DA-seq MSC clusters generated in Figure 3C. Corresponding Seurat scores were visualized on violin plots. Letters indicate statistically different conditions as defined by Student t-test with p value adjustment by Holms correction.

#### Supplemental Figure S5.

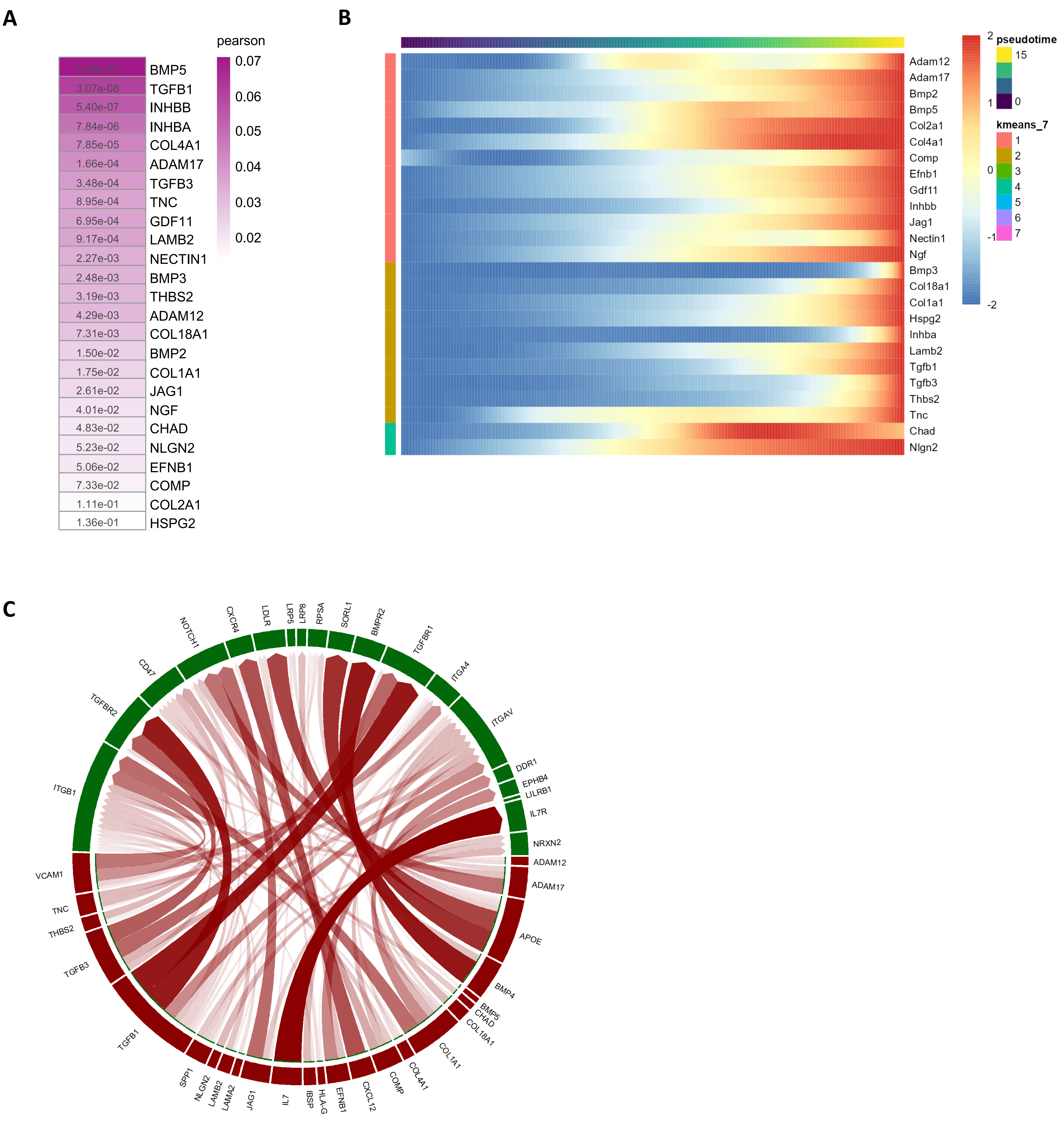


**Supplemental Figure S5. Impact of BM stromal cells on lymphoma B cells (related to Figure 5).**

(A) Heatmap representation of NicheNet-predicted interactions using B-cell-primed MSCs as senders and DOHH2 D40 cells as receivers. Color scale corresponds to pearson coefficient (quantifying ligand activities). The p-values assessed by random permutation were indicated in the heatmap.

(B) Expression profile along the pseudotime of the Top-25 ligands expressed by LepR$^{pos}$ MSCs.

(C) Circos plot showing predicted interactions between LepR$^{pos}$ MSCs and lymphoma B cells. MSC_Gr.Late were defined as senders and DOHH2 D40 as receivers.

### Supplemental Figure S6.

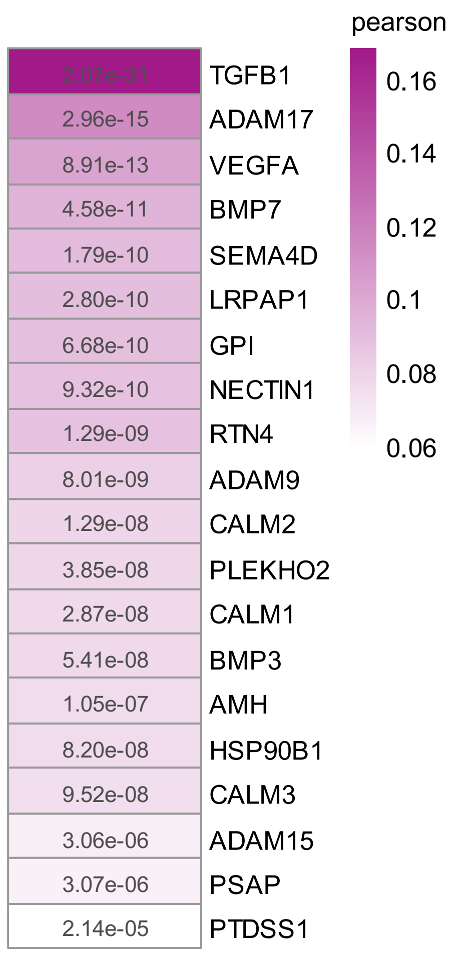


**Supplemental Figure S6. Impact of B cells on BM stromal cells (related to Figure 6)**

Heatmap representation of NicheNet-predicted interactions using DOHH2 D40 as senders and B-cell-primed MSCs as receivers. In boxes, colors correspond to pearson coefficient (quantifying ligand activities) and text to p value assessed by random permutation.

### Supplemental Figure S7.

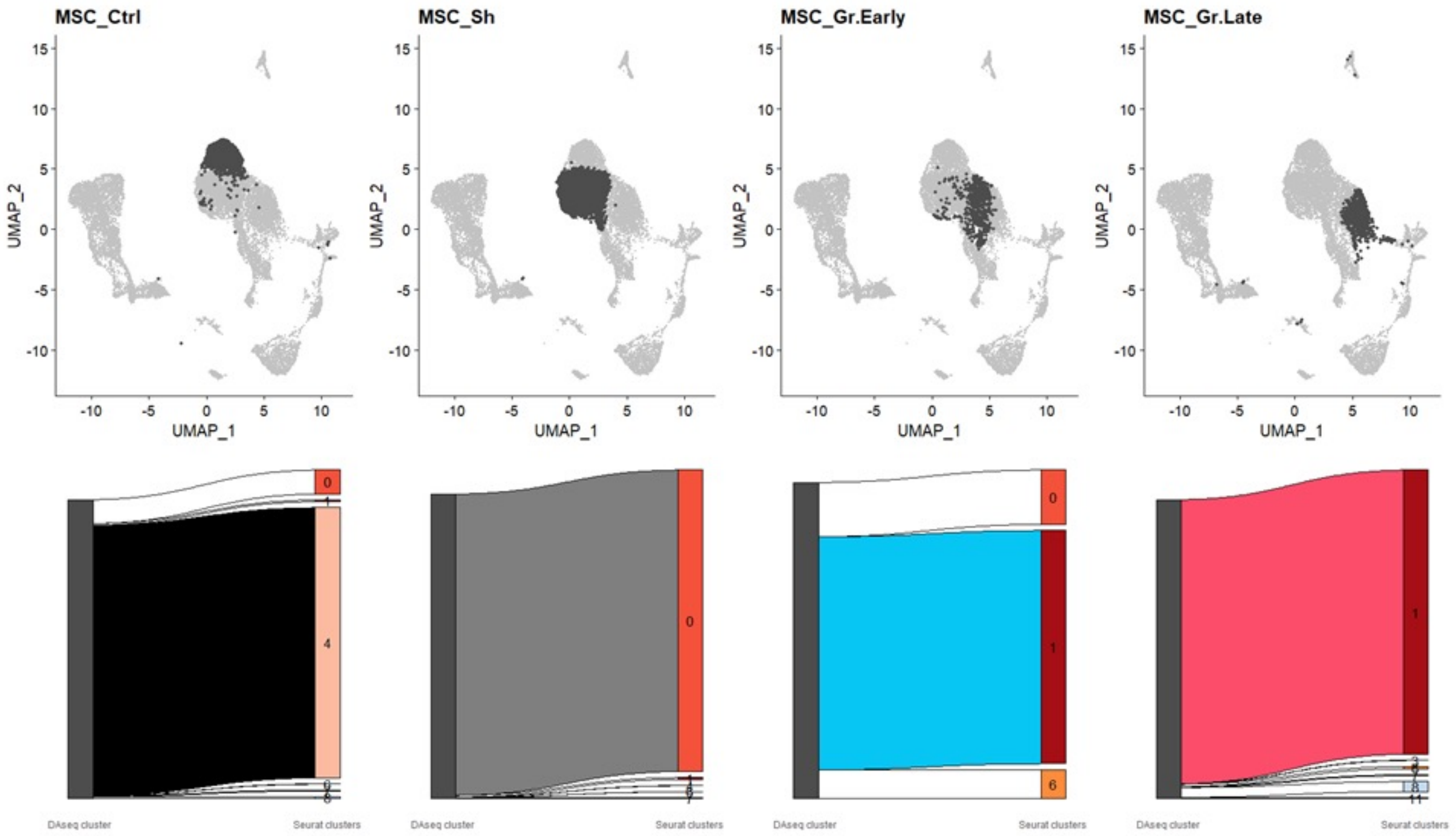


**Supplemental Figure S7. Relationship between DA-seq and Seurat clusters**

(Upper panel) Each unfiltered DA-seq-cluster was plotted on the UMAP plot of the scRNAseq analysis.

(Lower panel) Each DA-seq cluster was filtered to exclude outlier cells not belonging to the corresponding unsupervised Seurat cluster.
